# Individual-based modelling of shelled pteropods

**DOI:** 10.1101/2021.10.21.465291

**Authors:** Urs Hofmann Elizondo, Meike Vogt

**Author notes:** (Urs Hofmann Elizondo). Email addresses:* (Meike Vogt).

## Abstract

Shelled pteropods are cosmopolitan, free-swimming organisms of biogeochemical and commercial importance. They are widely used as sentinel species for the overall response of marine ecosystems to environmental stressors associated with climate change and changes in ocean chemistry. However, currently we are unable to project the effects of climate change on shelled pteropods at the population level, due to the missing spatio-temporal characterization of the response of pteropods to environmental stressors, and the limited information on the pteropod life history and life cycle. In this study, we implement a shelled pteropod Individual-Based Model (IBM), i.e. we simulate a pteropod population as a set of discrete individuals over several generations, life stages (eggs, larvae, juveniles and adults) and as a function of temperature, food availability and aragonite saturation state. The model is able to provide an abundance signal that is consistent with the abundance signal measured in the temperate region. In addition, the modeled life stage progression matches the reported size spectrum across the year, with two major spawning periods in spring and fall, and maturation in March and September. Furthermore, our IBM correctly predicts the abundance maxima of younger, smaller and potentially more susceptible life stages in spring and winter. Thus, our model provides a tool for advancing our understanding of the response of pteropod populations to future environmental changes.

## 1. Introduction

Shelled pteropods or sea butterflies, are cosmopolitan marine sea snails that spend their entire life cycle as free-swimming organisms (Bednaršek et al., 2012a; Suárez-Morales et al., 2009). Pteropods represent a major food source for other zooplankton, seabirds, whales, and fish (Falk-Petersen et al., 2001; Hunt et al., 2008), including commercially important ones such as herring, mackerel, or salmon (LeBrasseur, 1966; Lalli and Gilmer, 1989). In addition, pteropods likely dominate the magnitude of carbon export among pelagic marine calcifiers (Buitenhuis et al., 2019). Pteropods act as a vector for the export of particulate organic carbon to deeper waters due to the ballasting effect of their shells (Klaas and Archer, 2002), they repackage buoyant particles into faecal pellets (Tréguer et al., 2003; Manno et al., 2009), and they produce mucus webs during feeding (Gilmer, 1972), which aggregate and trap, and sediment particles that would not sink otherwise (Noji et al., 1997).

Pteropods represent an ideal sentinel species for the response of marine ecosystems to ocean acidification due to their soluble aragonite shells that are sensitive to the state of the ocean carbonate chemistry. (Orr et al., 2005; Bednaršek et al., 2017b). Pteropods produce thin shells made out of the metastable calcium carbonate mineral aragonite (Lalli and Gilmer, 1989), which is approximately 50% more soluble than its calcite counterpart (Mucci, 1983). Thus, a decrease in seawater carbonate concentration and pH due to the marine uptake of atmospheric *CO*_2_ (Doney et al., 2009) has been observed to lead to widespread pteropod shell dissolution and a decrease in their survival (Wall-Palmer et al., 2013; Bednaršek et al., 2017b; Manno et al., 2017). In addition, the exposure to lower carbonate concentration and pH decreases shell growth (Comeau et al., 2010b; Bednaršek et al., 2014), calcification (Moya et al., 2016), egg organogenesis (Manno et al., 2016), downward swimming velocities (Bednaršek et al., 2014; Bergan et al., 2017), and mechanical protection against predators (Lalli and Gilmer, 1989; Comeau et al., 2010a). In addition, the exposure to lower carbonate concentration is linked to developmental delays (Thabet et al., 2015), shell malformations or complete loss of shells (Comeau et al., 2010a). The exposure to corrosive waters not only affects pteropods, but has far reaching consequences for biogeochemical cycles (e.g. Bednaršek et al., 2014) and the flow of energy across trophic levels and from surface to depth (e.g. Capuzzo et al., 2017). For instance, developmental delays and impaired growth rates reduce the carbon export to deeper layers due to a diminished ballasting effect of pteropod shells (Klaas and Archer, 2002; Bednaršek et al., 2014), or decrease the flow of energy and carbon across trophic levels (Capuzzo et al., 2017), which diminishes fisheries yield (Ware, 2005) and global food resources (Chassot et al., 2010). The impacts of ocean acidification on the survival of pteropods depend not only on the magnitude and duration of the exposure to corrosive waters, but on the life stage of the pteropods (Bednaršek et al., 2016), as well as their exposure history (Bednaršek et al., 2017a).

Although pteropods are used as sentinel species for ocean acidification (Orr et al., 2005; Bednaršek et al., 2017b), available in-situ time series studies, while having the highest degree of realism and ability to quantify population-level responses to acidification (Andersson et al., 2015), have shown no change (Ohman et al., 2009; Head and Pepin, 2010; Thibodeau et al., 2018), declining (Beaugrand et al., 2012; Beare et al., 2013; Mackas and Galbraith, 2011), and increasing (Howes et al., 2015) trends in pteropod abundance around the world. This lack of relationship between pteropod abundance and acidification is likely linked to the variable responses of pteropods in nature (Bednaršek et al., 2016), or that the tipping point at which ocean acidification influences pteropods on a population level has not been crossed (Mackas and Galbraith, 2011; Doo et al., 2020). However, the latter stands in contrast to the observed widespread shell dissolution (Bednaršek and Ohman, 2015), suggesting that studies of abundance time series might have neglected indirect or sublethal effects that leave the abundance unaffected but influence shell structure, morphology, vertical distribution, or population demographics (Ohman et al., 2009; Bednaršek and Ohman, 2015; Manno et al., 2017; Bednaršek et al., 2019; Doo et al., 2020). Thus, a spatio-temporal characterization of shell dissolution that includes the life cycle of pteropods over several generations, with life stage specific responses and sensitivities to multiple stressors (e.g. ocean acidification and heat waves) will likely provide an accurate picture of the effects of acidification on pteropod populations (Johnson et al., 2016; Bednaršek et al., 2016; Manno et al., 2017; Bednaršek et al., 2019).

One possible approach to resolve the aforementioned limited spatio-temporal characterization of shell dissolution in pteropod populations is the simulation of pteropod populations as a set of discrete individuals over several generations using an Individual-Based Model (IBM; DeAngelis and Grimm, 2014). This approach requires a generalized life cycle model of shelled pteropods and their growth rates. Despite the ubiquity, commercial and biogeochemical importance of pteropods, their life cycle and growth rates have not been fully described (Hunt et al., 2008; Manno et al., 2017), due to the difficulty to culture them in a laboratory (Howes et al., 2014), and the dependence of their life cycle and longevity on local environmental conditions and taxonomic affiliation (Bednaršek et al., 2012a; Wang et al., 2017; Manno et al., 2017). However, unlike pteropods in the polar regions (e.g. Kobayashi, 1974; Gannefors et al., 2005; Hunt et al., 2008; Bednaršek et al., 2012b), a life cycle with two generations has been reported across different regions and species in the temperate zone, such as in the east coast of South America for *L. retroversa australis* (Dadon and de Cidre, 1992), the coast of Nova Scotia and the Gulf of Maine for *L. retroversa* (Lalli and Gilmer, 1989; Maas et al., 2020), or the temperate North Pacific for *L. helicina* (Wang et al., 2017). The life cycle of temperate pteropods has an overwintering generation that grows slowly during winter, reaching a size between 1 mm and 3 mm by spring, spawns the spring generation, and dies shortly after (Dadon and de Cidre, 1992; Wang et al., 2017; Maas et al., 2020). The spring generation grows rapidly to maturity, reaches relatively smaller sizes (0.5 mm - 1 mm) and dies shortly after spawning the next overwintering generation in late autumn (Dadon and de Cidre, 1992; Wang et al., 2017; Maas et al., 2020). The life cycle and growth rates of pteropods in temperate regions are likely less influenced by seasonal changes in environmental conditions compared to their polar analogue species (Manno et al., 2017), and the longevity and growth rates of temperate species are consistent across species and regions. Additionally, recent culturing experiments of the temperate *L. retroversa’s* life cycle identify ten stages and the developmental timings of key organs/traits (Howes et al., 2014; Thabet et al., 2015), which refines the description of the life cycle of temperate pteropods, and thus allows for the modelling of specific effects of multiple stressors at the organismal to the population level.

The different life stages in the pteropod life cycle are separated by the development of key organs (Thabet et al., 2015). The development of key organs includes the larval shell (protoconch), wings (parapodia), and female gonads after six days, three weeks, and three months from hatching, respectively (Thabet et al., 2015). This detailed life cycle description with the developmental timings of key organs is crucial for the interpretation of the response of pteropods to ocean acidification, since the development of the protoconch marks the onset of early shell dissolution (Thabet et al., 2015; Johnson et al., 2016), which likely diminishes the ability for pteropods to deal with upcoming shell dissolution (Bednaršek et al., 2017a), or the development of parapodia marks the onset of the daily upward and downward migration of pteropods across the water column, i.e. diel vertical migration (DVM; Lalli and Gilmer, 1989; Mackas et al., 2005; Hunt et al., 2008), which exposes pteropods to corrosive waters at deeper layers and non-corrosive ones near the surface (Bednaršek and Ohman, 2015). An additional size or stage dependent trait is the calcification rate or shell repair rate (Comeau et al., 2010b), which increases with pteropod size (Bednaršek et al., 2014), making juveniles performing DVM relatively more susceptible to shell dissolution than adult organisms. However, the shell repair process is debated in the literature, with Mekkes et al. (2021) finding that shelled pteropods may produce thinner shells as a response to increased acidification. Given the consensus in the life cycle of temperate pteropods across regions and species (Lalli and Gilmer, 1989; Dadon and de Cidre, 1992; Wang et al., 2017; Maas et al., 2020), and the recent success in culturing experiments detailing the stages and developmental timings (Howes et al., 2014; Thabet et al., 2015), a pteropod IBM can be implemented based on the life cycle of temperate pteropods.

IBM’s have been used successfully in marine applications to characterize the life history of planktonic organisms, or life stages and the interaction between individuals and their surroundings (e.g. Werner et al., 1997; Parada et al., 2003; Miller et al., 1998; Dorman et al., 2015a). IBMs are well suited to link parameterised responses of individuals to environmental drivers and predict the overall responses of populations (Dorman et al., 2011; Stillman et al., 2014). In IBMs the population-level response is the sum over all individuals and their interactions between each other and their environment (DeAngelis and Grimm, 2014; Dorman et al., 2015b), and they are thus able to consider variations among individuals of a population, and along their life cycle (DeAngelis and Grimm, 2014). This accounts for the dynamic and heterogeneous environmental conditions that determine population-level characteristics, such as size or life stage structure, in time and space (Stillman et al., 2014). In the case of shelled pteropods, their response to ocean acidification has been shown to depend on their life stage (Bednaršek et al., 2016), life history (Bednaršek et al., 2017a), behaviour (Bednaršek and Ohman, 2015) and size (Bednaršek et al., 2014). For instance, younger pteropods are more sensitive to corrosive waters relative to adult pteropods (Bednaršek et al., 2016), smaller organisms are predominantly found near the surface, while larger ones might actively avoid corrosive waters (Bednaršek and Ohman, 2015), or the shell repair rates (Bednaršek et al., 2014), allow individuals to compensate from the shell damage due to corrosive conditions (Comeau et al., 2010b). Thus, a pteropod IBM considers indirect or sublethal effects of acidification, and unlike traditional time series analyses, might be able to aid in finding population-level responses to acidification at different time scales, from ocean acidification extremes to climate change trends.

In this study, we implement a shelled pteropod IBM using the life cycle with two generations of temperate pteropods (Lalli and Gilmer, 1989; Dadon and de Cidre, 1992; Wang et al., 2017; Maas et al., 2020), the developmental timings of their life stages (Howes et al., 2014; Thabet et al., 2015) and a temperature and food availability dependence on shell growth (Buitenhuis et al., 2019). In addition, we implement the acidification sensitivity along the life cycle of pteropods (Bednaršek et al., 2016), calcification rate with changing shell size (Bednaršek et al., 2014), exposure to acidification due to a size dependent DVM (Maas et al., 2012; Bednaršek and Ohman, 2015), and swimming velocities (Bergan et al., 2017), to analyze the outcome of our IBM on the output of a circulation model for the California Current System (CalCS). This allows us to test the hypotheses that the pteropod IBM (i) reproduces the pteropod abundance signal measured in the temperate region, (ii) reproduces the measured size spectrum across seasons, and (iii) characterizes the spatio-temporal distribution of pteropod life stages in the CalCS.

## 2. Model description

The description of the model follows the ODD (Overview, Design concepts, Details) protocol for IBMs (Grimm et al., 2006, 2010, 2020). The IBM is implemented in Python 3.7, and is available in the electronic supplementary material.

### 2.1. Purpose and patterns

The first purpose of this model is to predict the number, size spectrum and population dynamics of shelled pteropods, and their spatio-temporal distribution in the CalCS. Ultimately, the purpose of this model, which will be presented in follow-up work, is to explore the relationship between pteropod populations and environmental stressors such as acidification across a range of different time scales. To this end, we impose a fixed life cycle with two generations for temperate pteropods (Wang et al., 2017) and developmental timings of life stages and key traits and behaviours (e.g. protoconch, parapodia, onset of diel vertical migration, and maturity; Howes et al., 2014; Thabet et al., 2015) as a function of shell size to simulate population dynamics. The impacts of stressors are implemented to reduce shell size growth, which allows our model to simulate the impacts of stressors on the life history, behaviour, population demographics and spatio-temporal distribution of individual pteropods.

### 2.2. Entities, attributes, and scales

Our model includes one entity, the shelled pteropods. The shelled pteropod are treated as a collection of individuals with a constant number of members, i.e. as *super-individual*, to improve the model speed and memory use (Scheffer et al., 1995). These pteropods are characterized by 15 attributes (Tab. 1). These include the ID, the *parent ID, parent size, generation, age, size, Egg Release Readiness (ERR) index, spawning events, water temperature experienced (T), aragonite undersaturation experienced (*Ω_*arag*_), the *chlorophyll-a concentration (chl), oxygen concentration (O*_2_*), accumulated damage (Dam_acc_), date* and *location*. The *ID* and *parent ID* are unique identifiers for each pteropod and the pteropod that spawned them. The *parent size* represents the shell length (mm) of the parent pteropod during spawning. The *generation* is an integer used to differentiate between pteropods of the spring (even integer) and overwintering (odd integer) generation as described for temperate regions (Dadon and de Cidre, 1992; Lalli and Gilmer, 1989; Wang et al., 2017; Maas et al., 2020). The *age* measures the life time of pteropods in days since spawning. The *size* represents the current shell size of the pteropods. The *ERR index* (similar to the Clutch Readiness Fraction presented in Miller et al., 1998), represents whether or not a mature pteropod is ready to spawn eggs. The *spawning events* represents the number of times a pteropod has produced and released eggs. The *T* and Ω_*arag*_ quantify the average water temperature in °C, and aragonite undersaturation experienced by a pteropod in one day, respectively. The 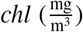 quantifies the cumulative sum of chlorophyll-a concentrations experienced by a pteropod (Wormuth, 1981; Hunt et al., 2008). The 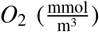 quantifies the oxygen concentration experienced by a pteropod throughout the day. The *Damacc* quantifies the amount of CaCO_3_ (mg) lost due to dissolution throughout the life of each pteropod after calcification and repair. The *date* and *location* depict the day in a year after the 1^*s*^*t* of January, and the depth (m), latitude (°*N*) and longitude (°*E*) of a pteropod. The *date* and *location* attributes enable us to analyze the IBM using a circulation model output. However, the IBM on its own does not have a spatial scale. The IBM’s total simulation time is not constrained, and is run at a one-day time-step.

**Table 1.** Summary of attributes characterizing the simulated shelled pteropods.

| Variable name | Variable type and units | Meaning |
| --- | --- | --- |
| ID | Static integer without units | Unique identifier for each pteropod |
| Parent ID | Static integer without units | Unique identifier of the parent pteropod |
| Parent size | Static real number in mm | Shell size of the parent pteropod at the time of spawning |
| Generation | Static integer without units | Integer used to identify whether a pteropod corresponds to the spring (even integer) or overwintering (odd integer) generation |
| Age | Dynamic integer in days | Number of days passed since the pteropod was spawned |
| Size | Dynamic real number in mm | Shell size or length of the pteropod |
| Egg Release Readiness (ERR) index | Dynamic real number between 0.0 and 1.0 without units | Fraction depicting whether a pteropod is ready to spawn eggs. Spawning only begins if $ERR \geq 1.0$. |
| Spawning events | Dynamic integer without units | Describes the number of times a pteropod has produced and released eggs. |
| Water temperature experienced ($T$) | Dynamic real number in $^{\circ}\text{C}$ | Average $T$ experienced by a pteropod in one day |
| Undersaturation experienced ($\Omega_{arag}$) | Dynamic real number without units | Average $\Omega_{arag}$ experienced by a pteropod in one day |
| Chlorophyll-a concentration experienced ($chl$) | Dynamic real number in $\frac{\text{mg}}{\text{m}^3}$ | Cumulative sum of $chl$ experienced by a pteropod in one day. |
| Oxygen concentration experienced ($O_2$) | Dynamic real number in $\frac{\text{mmol}}{\text{m}^3}$ | Oxygen concentration experiences by a pteropod throughout a day. |
| Accumulated damage ($Dam_{acc}$) | Dynamic real number in $\text{mg CaCO}_3$ | Net amount of $\text{CaCO}_3$ lost due to dissolution throughout the life of a pteropod |
| Date | Dynamic integer for the date | Current date |
| Location | Dynamic real numbers in m, $^{\circ}\text{N}$ and $^{\circ}\text{E}$ | Location of the pteropod given in depth, latitude and longitude |

### 2.3. Process overview and scheduling

In this section we provide an overview of the processes that occur in our IBM, the order in which they are executed, and focus on how the attributes are changed. Out of the 15 attributes, the *age, size, ERR index, spawning events*, *T*, Ω_*arag*_, *chl, O*_2_, *Dam_acc_, date and location* are changed on each one-day time-step by the following five processes (Figure 1): (1) Movement and environment tracker, (2) Mortality, (3) Shell growth, (4) Development, (5) Spawning eggs.

1. *Movement and environment tracker*: This function computes the vertical and horizontal movement of pteropods and tracks the environmental conditions, i.e. the *T*, *chl, O*_2_, and Ω_*arag*_ that pteropods experience on an one-hour time-step at each location. This higher temporal resolution was chosen, since vertical migration can result in exposure to corrosive and non-corrosive waters within a single day (Bednaršek and Ohman, 2015). In order to fit the simulation time-step of one day, we calculate the average *T*, Ω_*arag*_ and the cumulative sum of *chl* experienced by each pteropods in a 24 hour window. The *Movement and environment tracker* was calculated using the Lagrangian ocean analysis tool Parcels (Probably A Really Computationally Efficient Lagrangian Simulator) v2.1.3 (Delandmeter, 2019). This Lagrangian tool uses custom Parcels objects for the execution, and thus we have to translate the information for each pteropod stored as a numpy array (Oliphant, 2006) into the custom Parcels object back and forth before and after the execution of the *Movement and environment tracker* function.
2. *Mortality*: This function computes whether a pteropod dies or continues to live based on daily size-dependent mortality rates, the *age* of the pteropods, *generation*, dissolution, and *number of spawning events*.
3. *Shell growth*: This function determines the net shell growth, i.e. growth, including dissolution and repair, given the pteropod *size*, and the *T*, Ω_*arag*_ and *chl* experienced by the pteropod in the *Movement and environment tracker*.
4. *Development*: This function determines the life stage that each pteropod has reached based on their *size*. Different life stages are linked to the development of key traits/organs (e.g. protoconch, parapodia, juvenile/adult shell, maturity). These key traits/organs are used in other functions, e.g. the development of the protoconch and parapodia to mark the onset of shell dissolution and DVM, respectively.
5. *Spawning*: This function determines whether a pteropod has reached maturity and is ready to spawn eggs.

**Figure 1.**
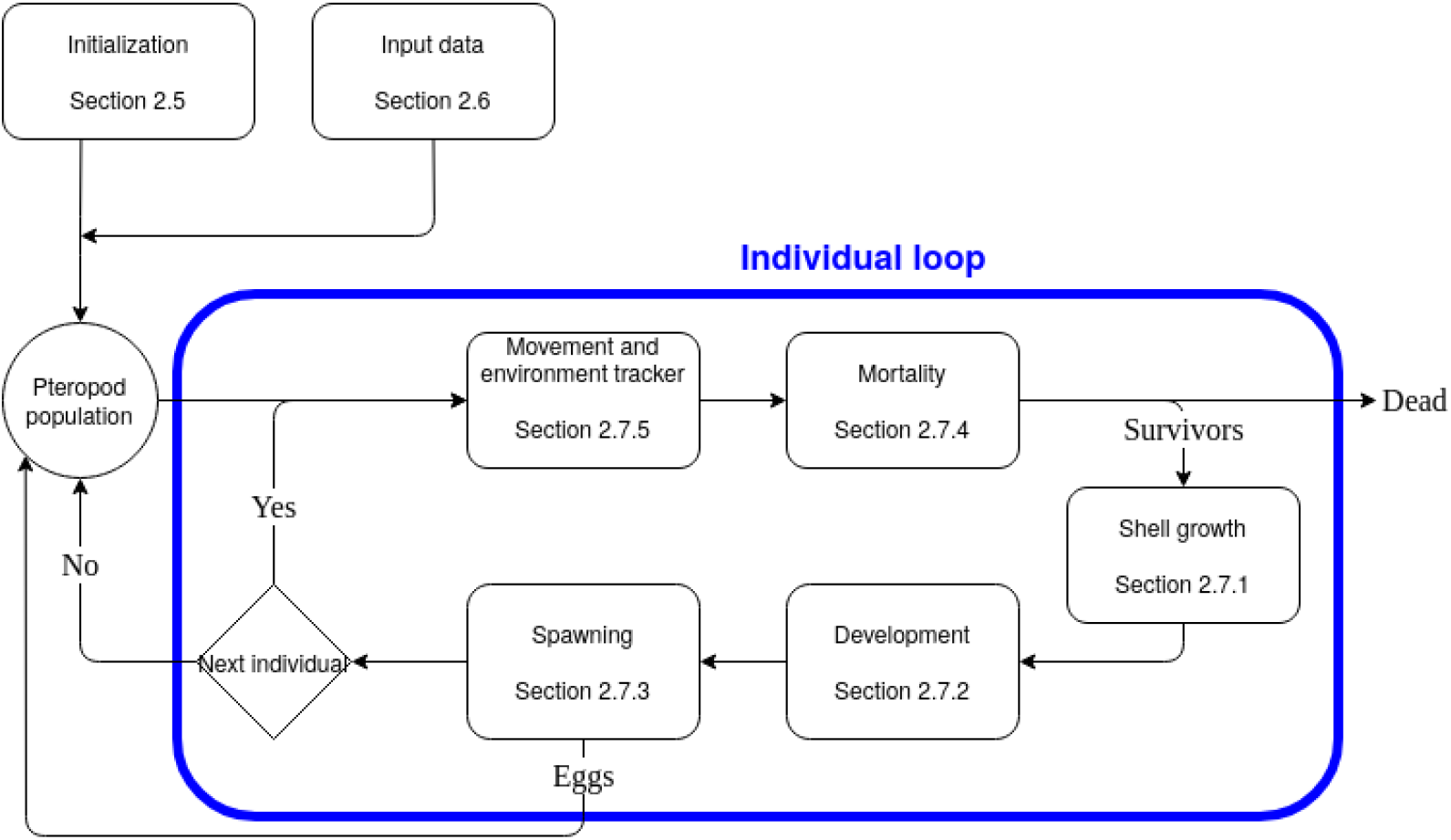
Pteropod individual-based model overview. The blue border shows the processes executed by each simulated pteropod on each one-day time-step.

**Figure 2.**
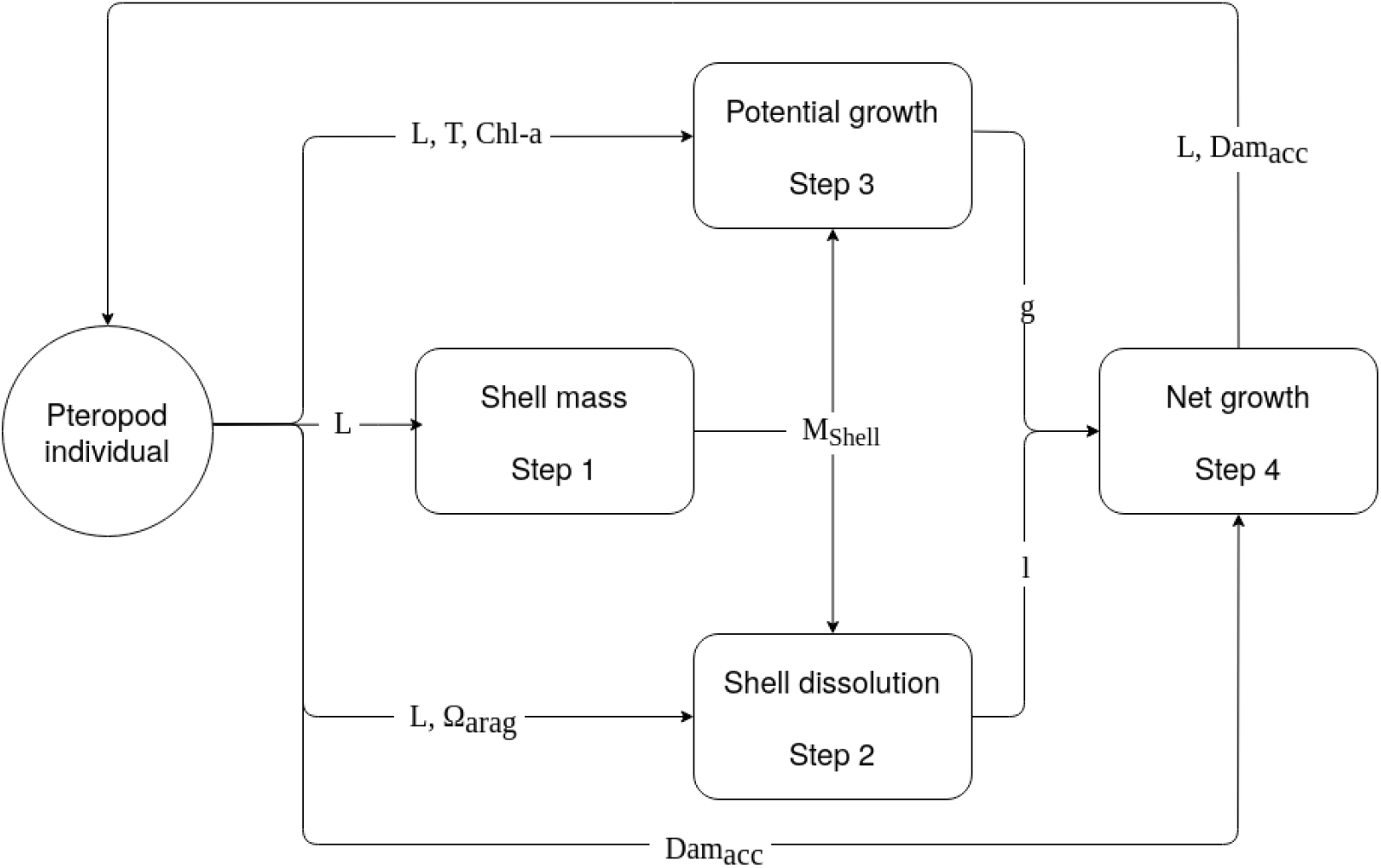
Structure of shell growth submodel divided into the four steps, and the attributes needed for each step. The shell mass step requires the pteropod size (L) and outputs the CaCO_3_ mass of the shell (*M_S hell_*). The potential growth requires L, *M_S hell_*, the temperature (T) and chlorophyll-a concentration (Chl-a) experienced by the pteropod and outputs the potential gain in shell mass *g*. The shell dissolution requires L, *M_S hell_*, and the aragonite saturation state (Ω_*arag*_) that the pteropod experienced, and outputs the loss in shell mass *l*. The net growth step requires the *g*, *l*, and damage accumulated (Dam_*acc*_) by the modeled pteropod. The final output of the net growth step is the updated L, and Dam_*acc*_.

### 2.4. Design concepts

#### Basic principles

In this paper, we implement a shelled pteropod IBM to characterize the life stage composition of pteropod populations in the CalCS. In accordance with previous findings, our model uses *size* as a key trait to determine the developmental timings of pteropods (Kiørboe and Hirst, 2014; Andersen et al., 2016). In addition, we link the growth and behaviour of individuals to environmental conditions such as temperature, Ω_*arag*_, food availability, and oxygen concentration based on observations and modelling efforts (e.g. Wang et al., 2017; Bednaršek et al., 2012b, 2014; Thabet et al., 2015; Buitenhuis et al., 2019).

#### Emergence

Pteropod population size structure, and developmental timings over several generations emerge from the temperature, food availability, size dependent sensitivity to acidification, repair rates, and the occurrence of undersaturated conditions in relation to their life cycle. As susceptible life stages may co-occur with more corrosive conditions, continuous exposure to corrosive conditions may result in developmental delays that ultimately shift the peak of pteropod abundances and population demographics in future generations.

#### Adaptation

Pteropods that have developed parapodia adapt their maximum DVM depth during their descent. Pteropods descend in the water column until they reach a maximum depth, which depends on their current size and the maximum depth at the current location. However, if the pteropods reach the upper boundary of the aphotic zone, they stop their descent prior to reaching their maximum DVM depth. In addition, individuals actively avoid hypoxic layers (<60 *μ*mol L^−1^; Keeling et al., 2010; Ekau et al., 2010) by swimming towards the surface. These behaviours are modeled as indirect objective seeking, i.e. individuals follow rules that have been observed in nature (Grimm et al., 2010, 2020).

#### Sensing

Pteropods in our model are assumed to sense their current depth, the light availability, oxygen, and the time at which the sun sets and rises in each day and location. Our modeled pteropods aim to reach a maximum depth during their descent, and upon reaching this depth the descent stops. The pteropods are assumed to perceive the sunrise and sunset time at their current location.

#### Stochasticity

Processes such as the onset of DVM and mortality, or egg release time after reaching maturity are not explicitly implemented in our model. However, we use a stochastic parameterisation to mimic the observed natural variability (Grimm et al., 2010, 2020), or to smooth the transition of one life stage to the next (Miller et al., 1998). With regard to the onset of DVM, the mechanism by which pteropods sense the sunrise and sunset times at each location is not modeled. Instead, the onset of the migration towards the surface and towards the deeper layers are determined by the timing of the sunset and sunrise at the current location and date of the pteropods (Bianchi and Mislan, 2015), respectively. Pteropod mortality in our model is represented using stochastic process formulations. First, we determine the survival probability of each pteropod based on their life stage. Second, we draw a pseudo-random number from a continuous uniform distribution over the interval [0, 1), where individual pteropods die if their survival probability is lower than the pseudo-random number. Finally, the time needed to release eggs after reaching maturity is determined by the *ERR index* (section 2.2). During initialization, the *ERR index* is drawn from a pseudo-random number from a continuous uniform distribution over the interval [−1, 1).

#### Observations

Data that can be collected from our IBM for testing and analysis include the abundance of pteropods, population life stage composition, size structure, and all 15 attributes throughout the simulation over time and space. In addition, individual level observations such as the shell size as a function of the experienced Ω_*arag*_, temperature, food availability, age, and generation can also be collected from our IBM.

### 2.5. Initialization

The aim of the initialization is two-fold. First, it eliminates effects of initial conditions by spinning up the model to a steady state such that initial conditions have a minimal effect on the model’s outcome (Grimm et al., 2020). Second, the model at steady state defines a starting pteropod population life stage composition for future simulation experiments (section 2.10). During the initialization, pteropod experienced idealized environmental conditions, and growth was not impaired by acidification. The idealized environmental conditions were taken from daily sea surface chlorophyll-a concentration and temperature climatologies between 30°N-60°N and 115°W-135°W. For chlorophyll-a, we used the Level-4 chlorophyll-a product from the EU Copernicus Marine Environment Monitoring Service (CMEMS, 2021). For temperature, we used the sea surface temperature from the World Ocean Atlas 2013 (Locarnini et al., 2013).

We ran the initialization for 5′000 time-steps, i.e. around 13 simulation years, starting with a population of 1′500 eggs of the spring generation, and the processes listed in section 2.3, except for the *Movement and environment tracker* function. By excluding the latter process, the exposure to acidification is turned off, i.e. pteropod growth is not impaired by stressors. The attributes of the initial 1′500 eggs are as follows: The IDs are set to a number between zero and 1′499, parent IDs and parent size to −1, generation, age and spawning events to zero, ERR to a random number in the interval [−1, 1), size to 0.15 *mm* (Wang et al., 2017), and the *Dam_acc_* to 0.0 mg CaCO_3_. The remaining attributes (*T*, Ω_*arag*_, *chl, O*_2_, date, and location) are not initialized at this point, since these attributes are only changed during the execution of the *Movement and environment tracker* function.

The outcome of the initialization, results in a pteropod population composition that is used for simulations under time-varying environmental conditions. In the following section we present the structure and source of the physical-biogeochemical forcing used to represent the time-varying environmental conditions.

### 2.6. Input data

In order to simulate changes in pteropod populations driven by environmental conditions, the pteropod IBM requires input from external sources to represent changes in environmental conditions over time. The input data for the physical-biogeochemical forcing of the IBM needs to be given for each grid cells with the attributes time in days since the start of the simulation, location given as depth (m), latitude (°N) and longitude (°E), maximum depth (m), land/water index, horizontal and vertical current velocities (u, v, and w in 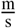), temperature (°C), Ω_*arag*_, chlorophyll-a concentration 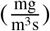, and oxygen concentration 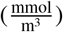 (Tab. 2).

**Table 2.**
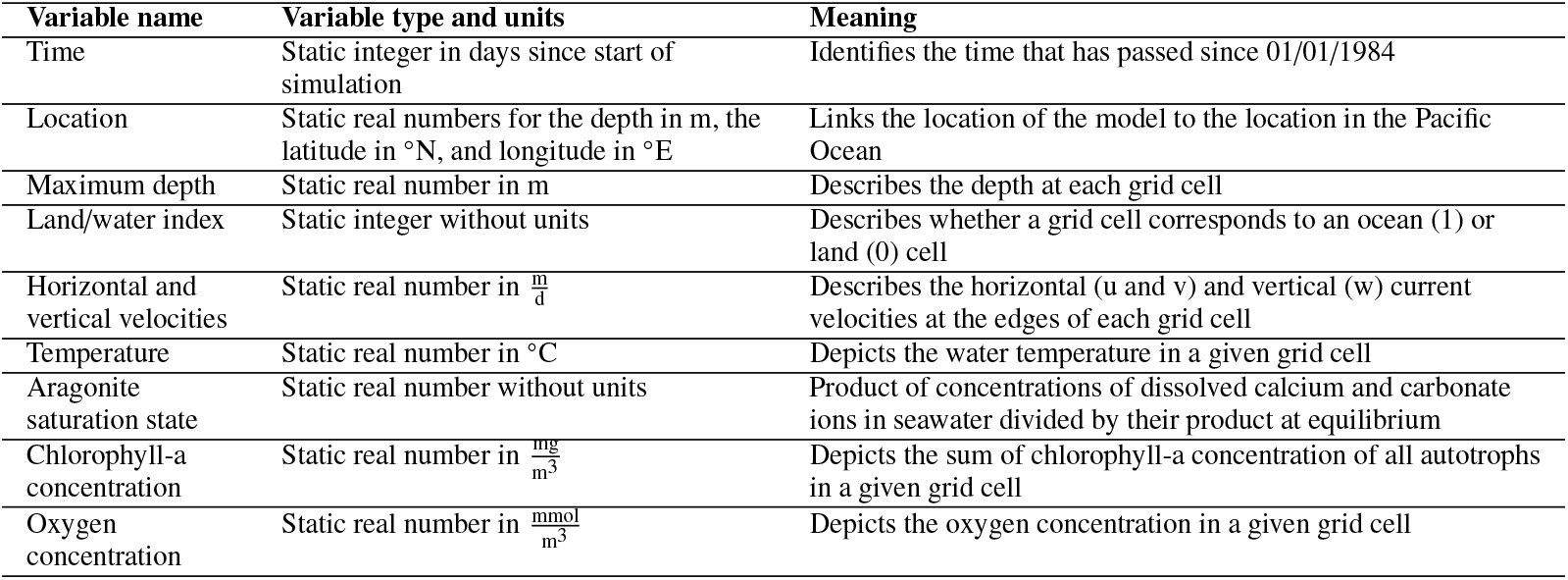
Description of the variables used to describe the grid cells.

| Variable name | Variable type and units | Meaning |
| --- | --- | --- |
| Time | Static integer in days since start of simulation | Identifies the time that has passed since 01/01/1984 |
| Location | Static real numbers for the depth in m, the latitude in °N, and longitude in °E | Links the location of the model to the location in the Pacific Ocean |
| Maximum depth | Static real number in m | Describes the depth at each grid cell |
| Land/water index | Static integer without units | Describes whether a grid cell corresponds to an ocean (1) or land (0) cell |
| Horizontal and vertical velocities | Static real number in $\frac{m}{d}$ | Describes the horizontal (u and v) and vertical (w) current velocities at the edges of each grid cell |
| Temperature | Static real number in °C | Depicts the water temperature in a given grid cell |
| Aragonite saturation state | Static real number without units | Product of concentrations of dissolved calcium and carbonate ions in seawater divided by their product at equilibrium |
| Chlorophyll-a concentration | Static real number in $\frac{mg}{m^3}$ | Depicts the sum of chlorophyll-a concentration of all autotrophs in a given grid cell |
| Oxygen concentration | Static real number in $\frac{mmol}{m^3}$ | Depicts the oxygen concentration in a given grid cell |

In this project we used the basin-wide hindcast simulation of Desmet et al. (in prep.) for the period between 01/01/1984 and 31/12/2019 to simulate the time-varying environmental conditions experienced by pteropods. This simulation has a daily average output of temperature, Ω_*arag*_, chlorophyll-a concentration, oxygen concentration, and horizontal and vertical current velocities modeled using the UCLA-ETH version of the Regional Oceanic Modeling System (ROMS; Marchesiello et al., 2002; Shchepetkin and McWilliams, 2005; Frischknecht et al., 2018) coupled with the Biological Elemental Cycling model (BEC; Moore et al., 2013; Frischknecht et al., 2017). The chlorophyll-a concentration was adjusted to the maximum and minimum chlorophyll-a concentrations found between 30°N-60°N and 115°W-125°W in the Level-4 chlorophyll-a product from the EU Copernicus Marine Environment Monitoring Service (CMEMS, 2021). The coupled model uses a horizontal curvilinear grid with its center at the US West Coast (Marchesiello et al., 2002; Frischknecht et al., 2018) and has 64 terrain-following vertical coordinates (Desmet et al., in prep.). This setup covers the Pacific Ocean basin with a horizontal grid spacing of 60 km that decreases to roughly 4 km towards central California (Frischknecht et al., 2018). The high resolution at the coast of California in the telescopic setup properly simulates upwelling conditions along the coastal region, and captures local (Frischknecht et al., 2018), as well as basin-wide processes (Frischknecht et al., 2015, 2017). The BEC model simulates the cycling of nutrients (e.g. carbon, phosphorus, nitrogen, silicon, iron), which govern the growth of a lower-trophic-level marine ecosystem represented by three phytoplankton functional types (small phytoplankton, diatoms, and diazotrophs) and one zooplankton type (Moore et al., 2013; Frischknecht et al., 2017). The Ω_*arag*_ is calculated offline using the MOCSY 2.0 routine (Dickson et al., 2007; Orr and Epitalon, 2015). In the following section we present in detail how the pteropods interact with the environment and how they develop.

### 2.7. Submodels

All model parameters and processes implemented in the IBM are summarized in the following sections. In our IBM, all five submodels (section 2.3) use *size* as an input, since it determines the developmental stage of the pteropods, their behaviour, and interaction with the environment. Thus, we first describe the *Shell growth* function, and build upon the assumption and parameterisations used in this function to describe the *Development*, *Spawning*, *Mortality*, and *Movement and environment tracker* functions. A summary of the functions and parameters are shown in Table A.4 and Table A.5.

#### 2.7.1. Shell growth

Measurements of the growth rate of shelled pteropods as a function of temperature and food availability are – unlike for other calcifying zooplankton (e.g. foraminifera; Lombard et al., 2009, 2011) – rare. In addition, measurements of pteropod growth rates are constrained to the Caribbean or the polar regions (see compilation in Bednaršek et al., 2016). To deal with these limitations, current models represent pteropods using the growth rates of better constrained and similarly sized zooplankton species (e.g. copepods Buitenhuis et al., 2019). However, a recent multi-year length-frequency analysis of pteropod populations parameterised the size of shelled pteropods as a function of time in the temperate North Pacific (Wang et al., 2017). This parameterisation provides a realistic progression of shell size throughout the life of a pteropod, and a growth rate that is specific to shelled pteropods. Thus, for our IBM, we define the size-specific growth rate as the derivative of the shell size parameterisation with respect to time and add key environmental drivers of growth, i.e. a temperature sensitivity based on the temperature coefficient *Q*_10_ (Buitenhuis et al., 2019), a food limitation term using the Monod equation (Monod, 1978; Lombard et al., 2011), and a shell dissolution function (Bednaršek et al., 2014). The implementation of these processes is described below.

Each one-day time-step, this function calculates the shell growth in four steps for each pteropod. First, we calculate the shell mass (*M_shell_*) in mg CaCO_3_. Second, we calculate the shell dissolution as a function of *M_shell_* and the Ω_*arag*_ that pteropods experienced. Third, we calculate the potential amount of CaCO_3_ in mg that pteropods can produce as a function of their current size, experienced temperature, and food availability. Finally, we calculate the growth or shell repair based on the potential amount of CaCO_3_ that can be produced and the amount that was lost due to dissolution.

##### Step 1 - Shell mass

We estimate the shell mass *M_shell_* in mg CaCO_3_ as a function of the *size L* as presented in Bednaršek et al. (2012b) and Bednaršek et al. (2014). The relationship between *M_shell_* and *L* is calculated using an empirical shell length-weight relationship derived from fitting an exponential function to shell length and dry weight (*DW*) measurements of *L. helicina* (Bednaršek et al., 2012b):

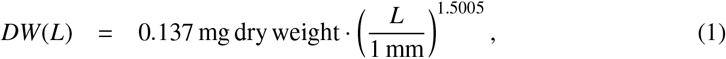

where *DW* is given in mg, and *L* is given in mm and non-dimensionalized. Next, we calculate *M_shell_* from *DW*. *DW* is transformed to total carbon content (*TC* in mg) using the average conversion factor 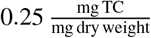 of dry weight to total carbon of pteropods (Davis and Wiebe, 1985). The total inorganic carbon (TIC) is calculated from *TC* based on the finding that 27% of the *TC* correspond to the *TIC* (Bednaršek et al., 2012b). *M_shell_* is calculated from *TIC* using the molecular mass ratio of C to CaCO_3_ 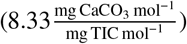. This results in the following relationship between *M_shell_* and *L*:

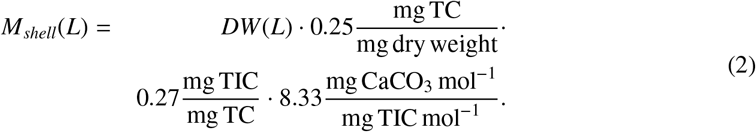

##### Step 2 - Shell dissolution

The damage to the shell as a function of the Ω_*arag*_ (*l* in mg CaCO_3_ d^−1^) is calculated based on the empirical fit of a two-parameter exponential function to measurements of the percentage pteropod shell loss per day (*v* in % shell loss d^−1^) as a function of Ω_*arag*_ in incubation experiments of *L. helicina* (Bednaršek et al., 2014):

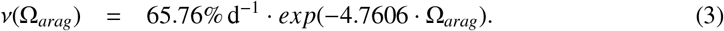

The exponential function *v* is able to capture shell mass loss at supersaturated conditions (Ω_*arag*_ > 1), and accurately describes the observed two-to three-fold increase in shell mass loss between Ω_*arag*_ = 1 and Ω_*arag*_ = 0.8 (Bednaršek et al., 2014). At higher supersaturation, the shell mass loss approaches zero (e.g. 0.052% at Ω_*arag*_ = 1.5, Figure A.8).

The *l* is then calculated by multiplying *M_shell_* by *v*:

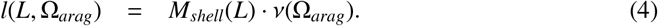

##### Step 3 - Potential growth

The amount of CaCO_3_ that pteropods can produce (*g* in mg CaCO_3_ d^−1^) is calculated as the amount of CaCO_3_ that a pteropod would produce after growth without dissolution based on their growth rate *μ*(*L, T, F*) as a function of size (*L*), temperature (*T*), and food availability (*F*). Explicitly, we subtract the shell mass calculated in step 1 (eq. 2) from the shell mass that pteropods would have after growth given their size, the temperature they experiences and food availability:

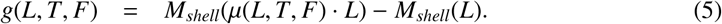

*μ* is a function of the current size *L*, the temperature that pteropods experience *T* (°C), and food availability 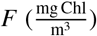. *μ*(*L, T, F*) consists of three terms, the reference growth rate *μ*_0_, the temperature sensitivity of growth Buitenhuis et al. (2019), and the food limitation of growth (Monod, 1978; Lombard et al., 2011):

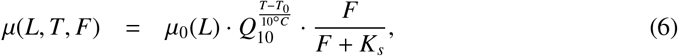

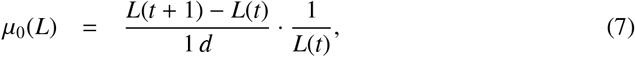

where *Q*_10_ = 1.3 is the unitless temperature coefficient (Buitenhuis et al., 2019), *T*_0_ = 14.5°C the reference temperature for growth (Wang et al., 2017), and *K_s_* the chlorophyll-a half saturation constant 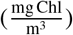. The value for 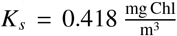 was determined based on our model tuning, which is explained in detailed in section 2.8.

The *μ*_0_, i.e. the size increase per day relative to the current size at *T*_0_, is calculated using the three-year cohort length-frequency analysis of *L. helicina* in the temperate North Pacific (Figure 3; Wang et al., 2017). The length-frequency analysis uses a variation of the *von Bertalanffy growth function* (von Bertalanffy, 1938), which considers seasonal changes in pteropod growth rates within one year (Wang et al., 2017), and is formulated as follows (Somers, 1988; García-Berthou et al., 2012; Pauly, 2013):

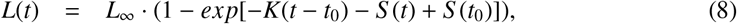

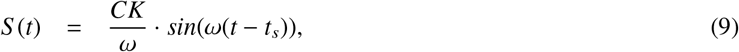

where *L*(*t*) is the size in mm at age *t*. *S* (*t*) parameterises the seasonal changes in pteropod growth. *ω* (in yr^−1^) is the recurrence rate of seasonal changes in pteropod growth. *t_s_* (in yr) marks the time sinusoidal growth oscillations begin with respect to a reference age *t*_0_. *L*_∞_ (in mm) denotes the maximum size of the pteropods, *K* (in yr^−1^) is the rate at which *L*_∞_ is approached, *t* is the age (given as fraction of a year), the reference age *t*_0_ (*yr*) is the age at which the pteropods have *L* = 0.0 mm. *C* is a unitless factor between zero and one that expresses the amplitude of growth oscillations. For this study, we use the values *t_s_* = −0.4 yr, *ω* = 2*π*yr^−1^ and *C* = 0.4 (Wang et al., 2017). The values for *K* = 5.07 yr^−1^ and *L*_∞_ = 4.53 mm are averaged from values reported in Wang et al. (2017) for the spring generation. The maximum life span of a generation is 300 days (Wang et al., 2017; Manno et al., 2017). As the size of pteropods is initialized as 0.15 mm (section 2.5; Wang et al., 2017), the values of *L*(*t*) are shifted upwards to match a starting position of 0.15 mm, i.e. *L*_0_ = 0.15 mm.

**Figure 3.**
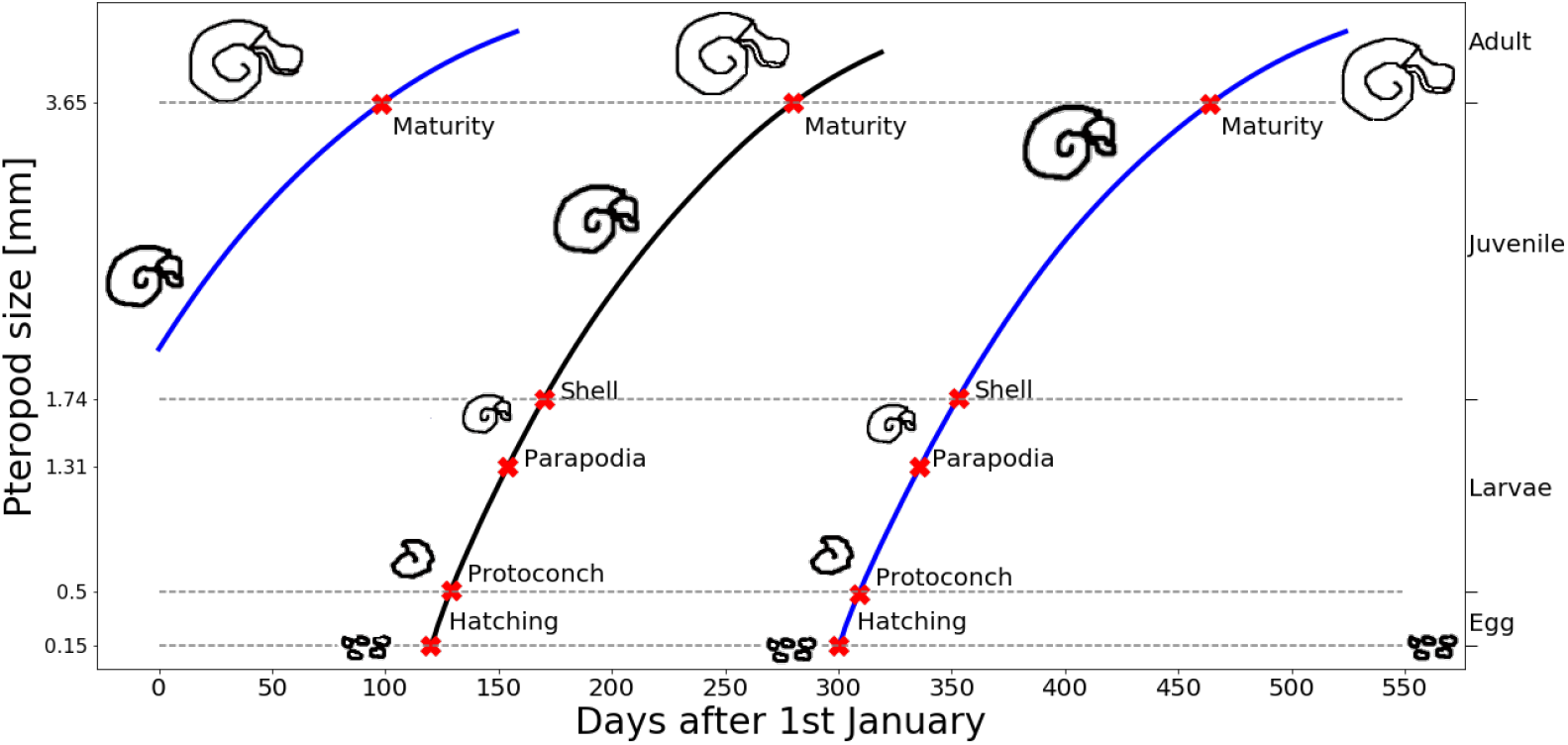
Size of pteropods parameterised as a function of time after hatching (Wang et al., 2017) for the spring (black) and winter (blue) generation. Annotated in red are the sizes at which pteropods hatch (0.15 mm), develop a protoconch (larval shell; 0.50 mm), parapodia (wings; 1.31 mm), juvenile/adult shell (1.74 mm) or reach maturity (3.65 mm) based on the comparison between culturing experiments (Howes et al., 2014; Thabet et al., 2015), and the parameterisation for the spring generation. The life stages (egg, larvae, juvenile, and adult) are shown on the right.

##### Step 4 - Net growth

We calculate the net amount of CaCO_3_ (*Net*_*CaCO*_3__) that remains after dissolution and calcification:

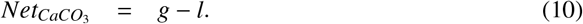

We then use *Net*_*CaCO*_3__. to calculate the effective change in shell mass Δ*M_shell_* and *Dam_acc_*:

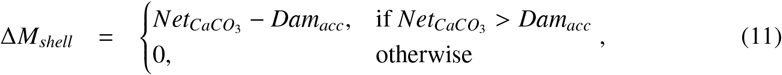

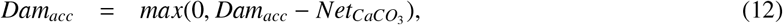

where an increase in shell mass only occurs if *Net*_*CaCO*_3__ is larger than the current *Dam_acc_*. Finally, to find the size change, we add Δ*M_shell_* to the *M_shell_*(*L*) calculated in step 1 (eq. 2) and extract the corresponding *L*.

#### 2.7.2. Development

Recent culturing experiments of the temperate *L. retroversa* have provided a detailed description of the life stage progression, and development of shelled pteropods (Howes et al., 2014; Thabet et al., 2015). The life cycle of *L. retroversa* includes the following stages (and developmental timings after spawning): the 2- (4 hours), 4- (6 hours), 8- (9 hours), 16-cell (11 hours), blastula (16 hours), gastrula (24-72 hours), hatching (3 days), trochophore (3-6 days), veliger (6-7 days), non-reproductive juveniles (1 month), and reproductive adult (3 months; Thabet et al., 2015). These life stages are linked to the development of key organs, including the larval shell (protoconch) and wings (parapodia) after six days, and three weeks from hatching, respectively (Thabet et al., 2015). This detailed life cycle description with the developmental timings of key organs is crucial for the interpretation of the response of pteropods to ocean acidification, since the development of the protoconch marks the onset of early shell dissolution (Thabet et al., 2015; Johnson et al., 2016), which likely diminishes the ability for pteropods to deal with upcoming shell dissolution (Bednaršek et al., 2017a), and the development of parapodia marks the onset of the daily upward and downward migration of pteropods across the water column, i.e. diel vertical migration (DVM; Lalli and Gilmer, 1989; Mackas et al., 2005; Hunt et al., 2008), which exposes pteropods to corrosive waters at deeper layers and non-corrosive ones near the surface (Bednaršek and Ohman, 2015).

The life stages that we diagnose in our IBM are based on the ten-stage life cycle model of *L. retroversa* (Howes et al., 2014; Thabet et al., 2015). However, we simplify the ten-stage life cycle into four life-stage tracers (Figure 3): the eggs (containing the 2-, 4-, 8-, 16-cell, blastula, gastrula, and trochophore stages), larvae (containing the veliger stage), juveniles (containing the non-reproductive juveniles), and reproductive adults. These four life stages were chosen based on the development of key organs. The transition from egg to larvae is based on the formation of the protoconch after six days of development (Thabet et al., 2015). During the larvae stage, pteropods develop parapodia after 21 days of development (Thabet et al., 2015). The transition from larvae to juvenile is based on the formation of the juvenile shell after 30 days of development (Thabet et al., 2015), and the transition from juvenile to adult is based on the development of female gonads after 90 days of development (Thabet et al., 2015). This definition facilitates the analysis of the model output by binning individuals that share physiological and behavioral characteristics (i.e. protoconch, parapodia, juvenile/adult shell, maturity) into categories.

In order to implement the four life stages in our IBM, we first defined size thresholds for the development of the key organs (protoconch, parapodia, juvenile shell, female gonads), using the pteropod size parameterisation (*L*(*t*), eq. (8), Figure 3). The protoconch develops at a size of *L*(*t* = 6 days) = 0.50mm, the parapodia at *L*(*t* = 21 days) = 1.31 mm, the juvenile shell at *L*(*t* = 30 days) = 1.74 mm, and the female gonads at *L*(*t* = 90 days) = 3.65 mm. On each time-step this function determines the life stage that a pteropod has reached depending on their size, and increases the *age* of the pteropods by one day.

#### 2.7.3. Spawning

The adults of pteropod populations in nature have synchronized widespread spawning events in spring and autumn (Lalli and Gilmer, 1989; Thabet et al., 2015; Wang et al., 2017), followed by a die-off of the largest individuals (Dadon and de Cidre, 1992; Wang et al., 2017; Maas et al., 2020). The eggs are released by adults as buoyant free-floating masses at the water surface (Lalli and Wells, 1978; Paranjape, 1968; Schalk, 1990; Gannefors et al., 2005; Comeau et al., 2010a; Manno et al., 2016). The number of eggs released per reproductive pteropod adult has a high variability, which is likely linked to the environmental conditions that pteropods experience (e.g. temperature, food availability, aragonite undersaturation; Paranjape, 1968; Manno et al., 2016). For instance, adult *L. helicina* release on average 5′936 eggs (observations range between 524 eggs and 10′051 eggs) in the Arctic (Lalli and Wells, 1978) and 565 eggs (observations range between 465 eggs and 708 eggs) in temperate regions (Paranjape, 1968). Across species and at the same region, the number of eggs released per adult can vary, where the number of eggs released by adult *L. retroversa* range between 83 eggs and 650 eggs in the Arctic (Lalli and Wells, 1978).

In our model we simulate the synchronized widespread spawning events in spring and autumn (Lalli and Gilmer, 1989; Thabet et al., 2015; Wang et al., 2017), the die-off of the largest individuals (Dadon and de Cidre, 1992; Wang et al., 2017; Maas et al., 2020), and neglect continuous low level spawning throughout the year (Lalli and Gilmer, 1989; Thabet et al., 2015; Wang et al., 2017). To this end, modeled adult pteropods only release eggs once, and die afterwards. We use a conservative estimate of 500 eggs per spawning event and per adult, since it is within the ranges reported for *L. retroversa* and *L. helicina* in the Arctic and temperate regions (Paranjape, 1968; Lalli and Wells, 1978). These eggs are released at the surface as free-floating masses, and the spawning time is spread in a 20 day time window after maturation using the *ERR* index (section 2.2). The latter indicates whether a modeled adult is ready to release eggs (*ERR* ≥ 1), and is used to smooth the shift from one life stage to the next (similar to the Clutch Readiness Fraction presented in Miller et al., 1998).

Each one-day time-step, we first increase the *ERR* index of the adults by 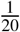. Second, we select the adults with *ERR* ≥ 1. Third, the selected adults spawn 500 eggs, where the attributes are initialized as follows: The unique IDs are determined based on the largest, ID of the population prior to spawning, parent IDs and parent size are the IDs of the parents and their current shell size, generation is set to the generation of the parent plus one, age and spawning events are set to zero, ERR to zero, size to 0.15 mm (Wang et al., 2017), and the *Dam_acc_* to 0.0 mg CaCO_3_. The attributes date, and location are initialized to the current date and at the surface at the current latitude and longitude of the parent. The remaining attributes (*T*, Ω_*arag*_, *chl, O*_2_) are initialized as missing values, as they are initialized during the execution of the *Movement and environment tracker* function on the next time-step. Finally, the attribute *spawning events* is set to one.

#### 2.7.4. Mortality

Information on the life span of temperate shelled pteropods is scarce (Manno et al., 2017). Under laboratory conditions, *L. retroversa* was recently shown to have a lifetime of six months (Howes et al., 2014; Thabet et al., 2015). In addition, the multi-year cohort analysis of Wang et al. (2017) indicates a lifetime of *L. helicina* of six months for a spring generation and eleven months for an overwintering generation (ten months on average), where a die-off of adult pteropods occurs shortly after the two major spawning events in spring and fall (Dadon and de Cidre, 1992; Wang et al., 2017; Maas et al., 2020).

In terms of daily mortality, information on daily mortality rates (*β*; Bednaršek et al., 2016) or mortality coefficients (M Lischka et al., 2011)) for different pteropod life stages is scarce, and reported values show large variability. For instance, juvenile *L. helicina* collected in Kongsfjord had a mean mortality coefficient of 12.6% (Lischka et al., 2011), whereas the mortality coefficient for juvenile *L. helicina* in the Scotia Sea is around 1.0% (Bednaršek et al., 2012b, 2016). However, the limited observations of pteropod mortality document the effect of the exposure to corrosive water to pteropod mortality for different undersaturation states and temperatures (Lischka et al., 2011), and thus a potential parameterisation for the increase in daily mortality as a function of the aragonite undersaturation state experienced by pteropods.

In our model, the pteropods die after reaching a maximum age, after releasing eggs, due to exposure to corrosive waters, and due to other causes. We chose a maximum age of ten months as averaged in the multi-year cohort analysis of Wang et al. (2017). We used the data reported in Lischka et al. (2011) to calculate the increase in mortality due to the exposure to corrosive waters. Due to the very limited information on *β* we tested 562′500 combinations of life stage and generation specific values for *β*. We chose the best combination of life stage and generation specific *β* that led to the best match-up between simulated pteropod abundance and observed pteropod abundance between 30°and 60°latitude in the Northern Hemisphere. The detailed tuning procedure is presented in section 2.8.

Each one-day time-step, we first calculate *M* for each life stage and generation based on the calibrated values for *β*. This means that we first calculate the proportion of individuals of each life stage that would die (Bednaršek et al., 2016):

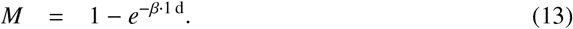

Second, for each modeled pteropod we draw a pseudo-random number from a discrete uniform distribution in the interval [0, 1), and compare this number for the life stage and generation specific *M*. If the pseudo-random number is smaller than *M* then the modeled individual dies. Third, we calculate the increase of *M* (Δ*M*) as a function of the Ω_*arag*_ experienced by each modeled pteropod (section Appendix A.2, Figure A.9) based on the observations of Lischka et al. (2011). Δ*M* is given by the difference between the undersaturation experienced by the modeled pteropod (Ω_*arag*_) and the reference saturation state where dissolution is negligible (Ω_*arag*_ = 1.5; section 2.7.1, step 2):

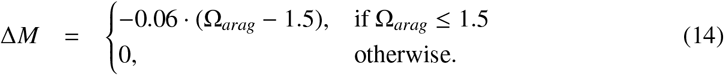

Fourth, we add Δ*M* to *M* and compare the pseudo-random number drawn at the beginning with the corrected mortality coefficient. If the pseudo-random number is smaller than *M* + Δ*M* then the modeled individual dies. Finally, we select the individuals with an age of ten months or those that have released eggs and remove them from the population.

#### 2.7.5. Movement and environment tracker

Shelled pteropods are able to remain neutrally buoyant for extended periods of time (e.g. during feeding; Lalli and Gilmer, 1989), and thus drift horizontally with the ocean currents. However, pteropods develop parapodia during their larvae stage (Thabet et al., 2015), which allow them to escape predation (Gilmer, 1972; Gilmer and Harbison, 1986), or swim closer to the surface at sunset to feed, and back to deeper waters to avoid predation at sunrise, i.e. perform diel vertical migration (DVM; Wormuth, 1981; Lalli and Gilmer, 1989; Mackas et al., 2005; Hunt et al., 2008; Maas et al., 2012; Bianchi and Mislan, 2015). Most shelled pteropods approximately remain in the upper 200 m of the water column (Lalli and Gilmer, 1989), where older and larger individuals tend to swim to deeper layers in comparison to younger and smaller individuals (Bednaršek and Ohman, 2015). Adult pteropods can reach upward swimming speeds of 44 mms^−1^, sinking speeds of 45mms^−1^ (Chang and Yen, 2012), and speeds between 80 mms^−1^ (*L. retroversa*) and 120 mm s^−1^ (*L. helicina*) during escape (Gilmer and Harbison, 1986). These relatively high swimming and sinking speeds imply that pteropods are able to swim several hundreds of meters in a short time span (one to two hours; Lalli and Gilmer, 1989), and thus pteropods can experience a wide range of environmental conditions in a single day, e.g. they can be exposed to corrosive conditions at deeper layers and non-corrosive conditions near the surface within a single day due to their vertical migration (Bednaršek and Ohman, 2015).

In our model, this function serves three purposes. First, we simulate the drift of the pteropods along the ocean currents, based on the horizontal and vertical current velocities calculated in the ROMS hindcast (section 2.6). To this end, we use the Lagrangian ocean analysis tool Parcels v2.1.3 (Delandmeter, 2019). Second and on top of the drift, we simulate the vertical migration of pteropods using size dependent swimming and sinking speeds (Chang and Yen, 2012), and a size dependent DVM depth (Bednaršek and Ohman, 2015). The onset of the upward migration towards the surface and downward migration towards the deeper layers are determined by the timing of the sunset and sunrise at the location and date of the pteropod (Bianchi and Mislan, 2015), respectively. Third, we calculate the average *T*, Ω_*arag*_, and *O*_2_ and the cumulative sum of *chl* that the pteropods experience throughout their movement in a day. To this end, the *Movement and environment tracker* function interpolates the physical-biogeochemical forcing from two daily average outputs (section 2.5) to then calculate the position of each pteropod and sample the environmental conditions that the pteropods experience throughout the day with a sub-timestep of one hour.

##### Current-driven drift

The current-driven drift of modeled pteropods is calculated using the horizontal and vertical current velocities of the ROMS hindcast simulation (section 2.6) using Parcels’ built-in fourth-order Runge-Kutta integration scheme including the vertical velocity (Delandmeter, 2019). This advection scheme has been reported to result in beaching of particles (Delandmeter, 2019). Thus, as recommended in Delandmeter (2019), we implemented an artificial current that pushes pteropods back to the ocean if they drift onto the shore, below the sea floor, or above the surface.

##### DVM

The depth of the modeled pteropods is only changed through DVM if they have developed parapodia, i.e. if modeled pteropods have a size of 1.31 mm (Figure 3). The vertical movement of modeled pteropods is constrained to the upper 250 m of the water column, since pteropods in the CalCS have mainly been observed to stay above this depth (Bednaršek and Ohman, 2015). The depth to which modeled pteropods sink (*d_DVM_*(*L*)) is calculated as the product of the maximum depth of 250 m and the ratio of the current pteropod size (*L*) to the maximum pteropod size (*L*_∞_ = 4.53 mm; section 2.7.1, step 3):

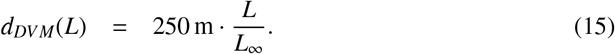

In cases where the water column is shallower than *d_DVM_*(*L*), modeled pteropods sink to the bottom of the water column.

After calculating *d_DVM_*(*L*), we determine the swimming and sinking velocities of the modeled pteropods based on their current size (*v*(*L*)). In analogy to the *d_DVM_*(*L*), we define the maximum swimming and sinking speed as 44 mm s^−1^ (Chang and Yen, 2012), and scale it by the ratio of *L* to *L*_∞_:

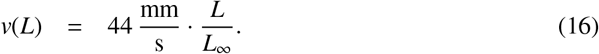

Our formulation is consistent with the description of Chang and Yen (2012), where the smallest modeled individuals (*L* = 1.30 mm) reach a swimming velocity of 12.6 mm s^−1^ and the largest ones *L* = 4.53 mm a velocity of 44 mm s^−1^.

Finally, to calculate the DVM, we simulate the amount of time that modeled pteropods spend at the surface, migrating between the surface and deeper waters, and at their DVM depth. To this end, we use the finding that zooplankton in general start swimming downwards around sunrise, and arrive at the surface from their maximum DVM depth around sunset (Bianchi and Mislan, 2015). The sunrise and sunset timings were calculated based on the date, and the position of each modeled pteropod (Nautical Almanac Office, 1990). Specifically, the onset of the downward migration (*t_down_*) corresponds to the sunrise time of the following day (*t_sunrise_*). *t_sunrise_* is determined for the following day, since the movement of pteropods is calculated for 24 hours starting at noon (section 2.6). For the onset of the upward migration (*t_up_*), we first determine the sunset time of the current day (*t_sunset_*) to obtain the time at which pteropods arrive at the surface. Next, we calculate the time at which the pteropods start their upward migration, as the time each pteropod would need to reach the surface from their maximum DVM depth given their size and their upward swimming speed (*v*):

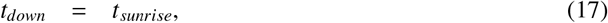

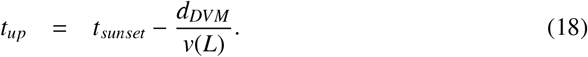

After reaching the surface, pteropods maintain their depth until *t_down_*. Similarly, pteropods that reach *d_DVM_*, maintain their depth.

##### Sampling of environmental conditions

The environmental conditions are provided by the ROMS hindcast simulation (section 2.6). On each one hour time-step of the *Movement and environment tracker* function (section 2.7.5), modeled pteropods sample their environment (temperature, chlorophyll-a concentration, oxygen concentration, and aragonite saturation state) at their location. After 24 one-hour time-steps, we calculate the average temperature, aragonite saturation state, chlorophyll-a concentration, and oxygen concentration that the pteropods experienced and update the attributes *T*, Ω_*arag*_, *chl*, and *O*_2_ with these values (section 2.3).

### 2.8. Model tuning

Information of the values of the half saturation constant for the food limitation *K_s_* (section 2.7.1) and the life stage and generation specific mortality rates (*β*; section 2.7.4) are limited or unknown. To find possible values for *K_s_* and *β* we performed a two-fold tuning exercise under idealized conditions (5′000 days; section 2.5). First, we estimate the value of *K_s_* to simulate the two generation per year cycle reported for temperate pteropods (Lalli and Gilmer, 1989; Dadon and de Cidre, 1992; Wang et al., 2017; Maas et al., 2020), i.e. the time span between a pteropod that hatches in spring (30th April) and the day that the next generation releases eggs is around 365 days. Second, we tested 562′500 combinations of mortality rates (*β*; section 2.7.4), and chose the best combination of life stage and generation specific *β* that led to pteropod populations with an abundance that best matches the observed pteropod abundance between 30°and 60°latitude in the Northern Hemisphere reported in the MARine Ecosystem biomass DATa (MAREDAT) initiative (henceforth referred to as the MAREDAT abundance; Bednaršek et al., 2012a) throughout a year.

With regard to *K_s_*, we tested values for *K_s_* ranging between 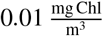 and 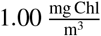. The duration of the two generation cycle increases linearly with increasing *K_s_* (Figure A.10), where 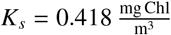 results in a two generation cycle of 365 days (Figure 3). This coupling of food availability and growth rate does not show a phase of slow growth during winter (overwintering) as presented in Wang et al. (2017). However, our approach is able to simulate phases of slow growth during periods of food scarcity followed by rapid growth in spring (Figure A.11).

Between and within pteropod life stages, the *β* can vary by more than a factor of ten (e.g. between eggs and all other life stages; Lischka et al., 2011; Bednaršek et al., 2016). Thus, for the life stage and generation specific *β* we tested the values (10, 20, 25, 35, 45, 50) · 10^−2^ d^−1^ for the eggs, and (1, 2.5, 5, 7.5, 10) · 10^−2^ d^−1^ for all other life stages. The combinations that resulted in stable population dynamics with at least 50 and at most 35’000 individuals (excluding eggs) were evaluated based on the similarity between the daily mean pteropod abundance across the last third of the simulation, and the daily pteropod abundance interpolated from the monthly abundance climatology observed in the MAREDAT abundance (Bednaršek et al., 2012a). The eggs modeled were not included to calculate the daily mean pteropod abundance, since the MAREDAT abundance does not contain egg counts. The similarity was measured based on the Pearson (*r*) and Spearman (*ρ*) correlation coefficient, the Manhattan distance (*L*1), and the proportion of modeled abundance (*f_out_*) outside of the abundance range (mean ± std. dev) reported in MAREDAT.

The values (10, 7.5, 1, 7.5) · 10^−2^ d^−1^ for the eggs, larvae, juvenile and adults of the spring generation and (45, 1, 1, 1) · 10^−2^ d^−1^ for eggs, larvae, juveniles and adults of the overwintering generation led to the most similar pteropod abundance pattern with *r* = 0.99, *ρ* = 0.91, *L*1 = 15.50, and *f_out_* = 0.29 (Figure 4). Differences between the MAREDAT abundance and the IBM simulation are found in the abundance magnitude in winter and the timing of abundance peaks in spring. The simulated abundance is mostly within the range reported in the MAREDAT abundance, except in winter, at the beginning of spring, and early summer. In addition, the spring pteropod abundance peak occurs four days later in the IBM than in the MAREDAT abundance.

**Figure 4.**
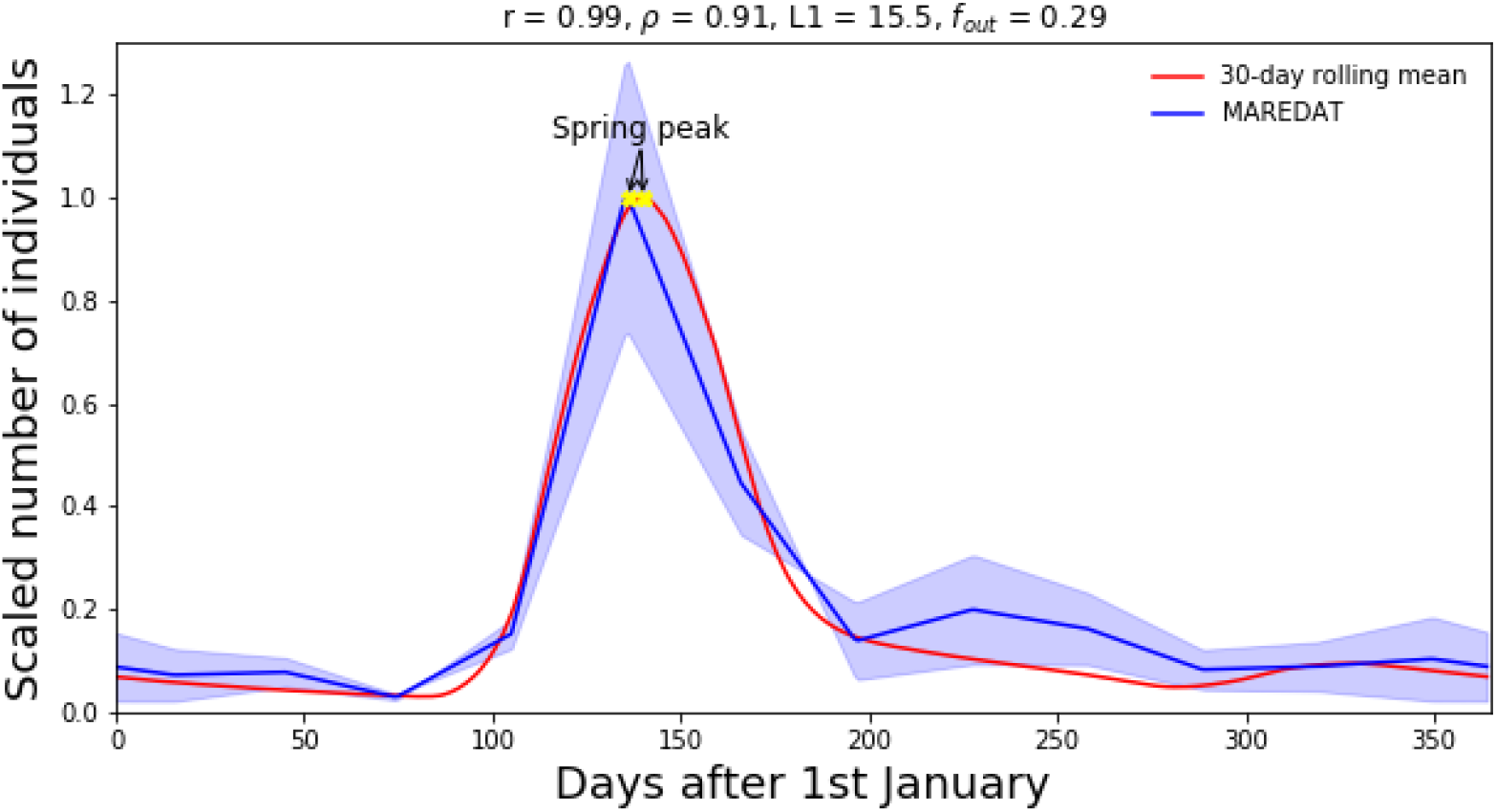
30-day rolling mean (red) of the daily pteropod abundance climatology simulated by the shelled pteropod IBM, and the daily abundance data and standard deviation of pteropods reported in the MAREDAT initiative (blue; Bednaršek et al., 2012a). The MAREDAT daily data was interpolated from monthly averaged and seasonally corrected abundance data found between 30°and 60°latitude in the Northern Hemisphere. The abundance data was scaled to the interval between zero and one. The spring pteropod abundance peak is shown in yellow. The Pearson correlation coefficient (*r*), Spearman correlation coefficient (*ρ*), the Manhattan distance (*L*1), and the proportion of modeled abundance outside of the measured abundance range (*f_out_*) between the MAREDAT data and the 30-day rolling mean are given at the top of the figure.

### 2.9. Sensitivity experiments

The daily mortality rates between and within pteropod life stages has been reported to vary by more than a factor of ten (section 2.7.3; Lischka et al., 2011; Bednaršek et al., 2016). Similarly, the range of the number of eggs that an adult pteropod releases can vary between 83 and 10′051 depending on the species and region (section 2.7.3; Lalli and Wells, 1978). These parameters represent a major uncertainty with regard to the life cycle of pteropods. Thus, we test the sensitivity of the IBM to the parameter choices associated with these two processes under idealized conditions (section 2.5). To this end, we increase and decrease the the number of eggs released by adults, the ERR, and each life stage and generation specific mortality rate by 10% individually. We evaluate the absolute abundance change in model-data-misfit based on the Manhattan distance (*L*1), and the seasonal changes caused by the increase/decrease in the chosen parameters using the Pearson correlation (*r*) to the pteropod abundance reported in MAREDAT (Bednaršek et al., 2012a).

Overall, the *L*1 between modeled pteropod abundance and the MAREDAT abundance increases for all sensitivity experiments, and *r* decreases by at most 0.12 (Table A.6). The largest difference in the modeled pteropod abundance is seen for the decrease by 10% of the number of eggs released per adult (Figure A.12I) with an increase in *L*1 of 27.1. The second largest change in modeled abundance is caused by an increase in the daily decrease in the mortality rate of the adults of the spring generation with *L*1 = 25.3 and *r* = 0.95 (Table A.6). This decrease in the parameter results in a longer spawning period, and thus a shift in the period of higher pteropod abundance relative to the unchanged simulation (Figure A.12G). For all other sensitivity runs, the modeled abundance pattern resembles the unchanged simulation albeit with mostly lower magnitude in the modeled abundance during spring and early summer (Figure A.12).

### 2.10. Modeling goals

In this study, we aim to (i) reproduce the pteropod abundance signal measured in the temperate region, (ii) reproduce the measured size spectrum across seasons, and (iii) characterize the spatio-temporal distribution of pteropod life stages in the CalCS. To this end, we run the shelled pteropod IBM for each year in the ROMS hindcast simulation (19984-2019) starting on the 1st of January until the 31st of December. The IBM is run as follows: First, we define an initial pteropod population based on the IBM initialization (section 2.5), which defines a starting pteropod population and life stage composition. Second, we randomly seed the initial pteropod population in the CalCS (Figure A.13) and initialize the depths and environmental conditions of the initial population using only the *Movement and environment tracker* function for three days. Third, we run the IBM for an entire year using all submodels (section 2.7). Thus, we start each simulation year with the same initial population. This approach was chosen, since it emphasizes the role of drift and environmental conditions on the pteropod population (e.g. Dorman et al., 2011, 2015a,b).

In order to compare the modeled pteropod abundance in the fully coupled model with the pteropod abundance signal measured in the temperate region, we calculate a pteropod abundance climatology based on the median from all IBM one-year runs for the entire hindcast period (01/01/1984 - 31/12/2019). We use this approach for the comparison, since the MAREDAT abundance data encompasses observations from several years (1951-2010; Bednaršek et al., 2012a). In addition, the climatology contains potential inter-annual variation in the modeled pteropod population, which might be more comparable to the MAREDAT abundance data. Next, we define the number of individuals within each *super-individual* (section 2.2), by dividing the maximum number of pteropods reported in the MAREDAT abundance across the region by the maximum number of modeled *super-individuals* in the climatology. Afterwards, we look at the pteropod population composition and the life stage progression from the climatology, and compare the life stage duration and developmental timings with observed life stage and developmental timings under laboratory conditions (Howes et al., 2014; Thabet et al., 2015), and qualitatively assess whether the modeled life stage progression and generation life span match field observations (Hsiao, 1939; Newman and Corey, 1984; Dadon and de Cidre, 1992; Lalli and Gilmer, 1989; Thabet et al., 2015; Wang et al., 2017; Maas et al., 2020). Lastly, we characterize the spatio-temporal distribution of pteropod life stages throughout a year based on their latitude and longitude. To this end, we determine the dominant life stage for each simulation day and 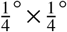 pixel. This approach allows us to characterize the dispersal and horizontal distribution of pteropod life stages during different developmental timings in the modeled pteropod life cycle.

## 3. Results

### 3.1. Population dynamics

The modeled pteropod community structure is characterized by a succession pattern of three different generations, which replace one another throughout the year (Figure 5). Across the year, the maximum median abundance of modeled *super-individuals* was 13′665.5 (excluding eggs, section 2.8), which is roughly 18 times lower than the maximum number of pteropods reported in the MAREDAT abundance across the region (249′960). Thus, each *super-individuals* represents a collection of 18 individuals. In general, the duration of both generations is similar in magnitude (Table 3). The spring generation is present in the population between 231 days (5th percentile) and 254 days (95th percentile). In comparison, individuals of the overwintering generation are present in the modeled population between 237 days and 271 days. The main difference between the spring and overwintering generation relates to their relative abundance. For instance, the spring generation reaches a maximum median abundance of 13′619.5 *super-individuals* (Figure 5A). In comparison, the larvae of the overwintering generation reach a maximum median abundance of 1′600 *super-individuals* (Figure 5B).

**Figure 5.**
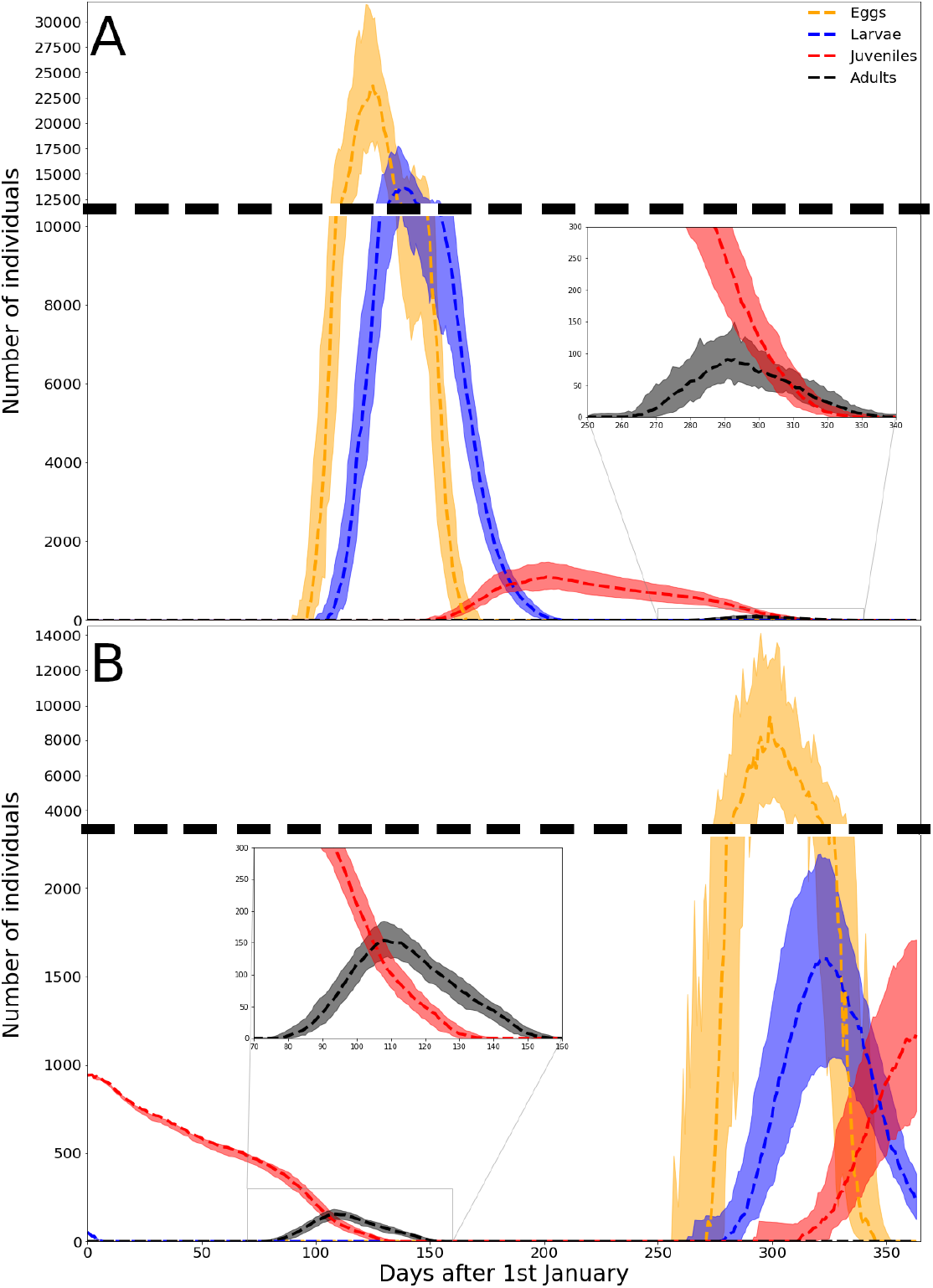
Abundance climatology of simulated pteropods of the (A) spring and (B) overwintering generation throughout the year in the California Current System. The shaded regions represent the 5th and 95th percentiles of all one year IBM runs. The abundance is divided into the life stages eggs (orange), larvae (blue), juveniles (red), and adults (black). The climatology was calculated from each one year IBM run on the ROMS hindcast for the period between 01/01/1984 and 31/12/2019.

**Table 3.** Climatological life stage succession pattern of the overwintering and spring shelled pteropod generations in the California Current System. The succession pattern was characterized based on the first appearance and disappearance of each life stage in each generation. The range in the timing of the succession events was calculated from all one year IBM runs.

|  |  | 5th percentile | 50th percentile | 95th percentile |
| --- | --- | --- | --- | --- |
|  |  | [Days after the 1st of January] |  |  |
| **Overwintering** | **Transition to larvae** | 287 | 278 | 263 |
|  | **Transition to juvenile** | 320 | 312 | 292 |
|  | **Transition to adult** | 81 | 77 | 72 |
|  | **Onset of spawning** | 100 | 97 | 90 |
|  | **Onset of die-off** | 152 | 157 | 164 |
| **Spring** | **Transition to larvae** | 110 | 106 | 100 |
|  | **Transition to juvenile** | 152 | 148 | 138 |
|  | **Transition to adult** | 270 | 262 | 248 |
|  | **Onset of spawning** | 279 | 272 | 257 |
|  | **Onset of die-off** | 331 | 337 | 344 |

Across all modeled years, the succession pattern of the pteropod generations and life stages followed the following sequence: Starting on the 1st of January, the juveniles of the overwintering generation start to mature and release eggs after 72-81 days and 90-100 days (95th and 5th percentile; Table 3). Shortly after spawning, the individuals of the overwintering generation start to die 152-164 days after the 1st of January (Table 3). The eggs of the spring generation begin their transition to the larval, juvenile and adult stage between 100-110, 138-152, and 248-270 days after the 1st of January (Table 3). Upon reaching maturity, adults of the spring generation begin to spawn the next overwintering generation at the earliest 257 days after the 1st of January and at the latest 279 days after the 1st of January (Table 3). The spring generation then disappears from the population 331-344 days after the 1st of January (Table 3). The eggs of the overwintering generation begin their transition to the larval and juvenile stage 263-287 days and 292-320 days after the 1st of January (Table 3), respectively.

Overall, the modeled life stage progression fits well with the reported pteropod life stage occurrence across multiple regions (Figure 6). The main difference between the modeled and observed life stage occurrence is seen for the modeled juveniles. The juveniles of the spring generation have been observed to occur between 171 and 264 days after the 1st of January (Newman and Corey, 1984; Dadon and de Cidre, 1992; Wang et al., 2017). However in our climatology, the juveniles of the spring generation are present between 148 and 327 days after the 1st of January (red in Figure 6). Similarly, the juveniles of the overwintering generation are present until 136 days after the 1st of January, which is 58 day longer than what has been observed (Dadon and de Cidre, 1992). In addition, the eggs of the overwintering generation, the larvae of the overwintering generation, and the adults of the spring generation are present 42, 17, and 33 days longer in the model compared to observations, respectively (Figure 6).

**Figure 6.**
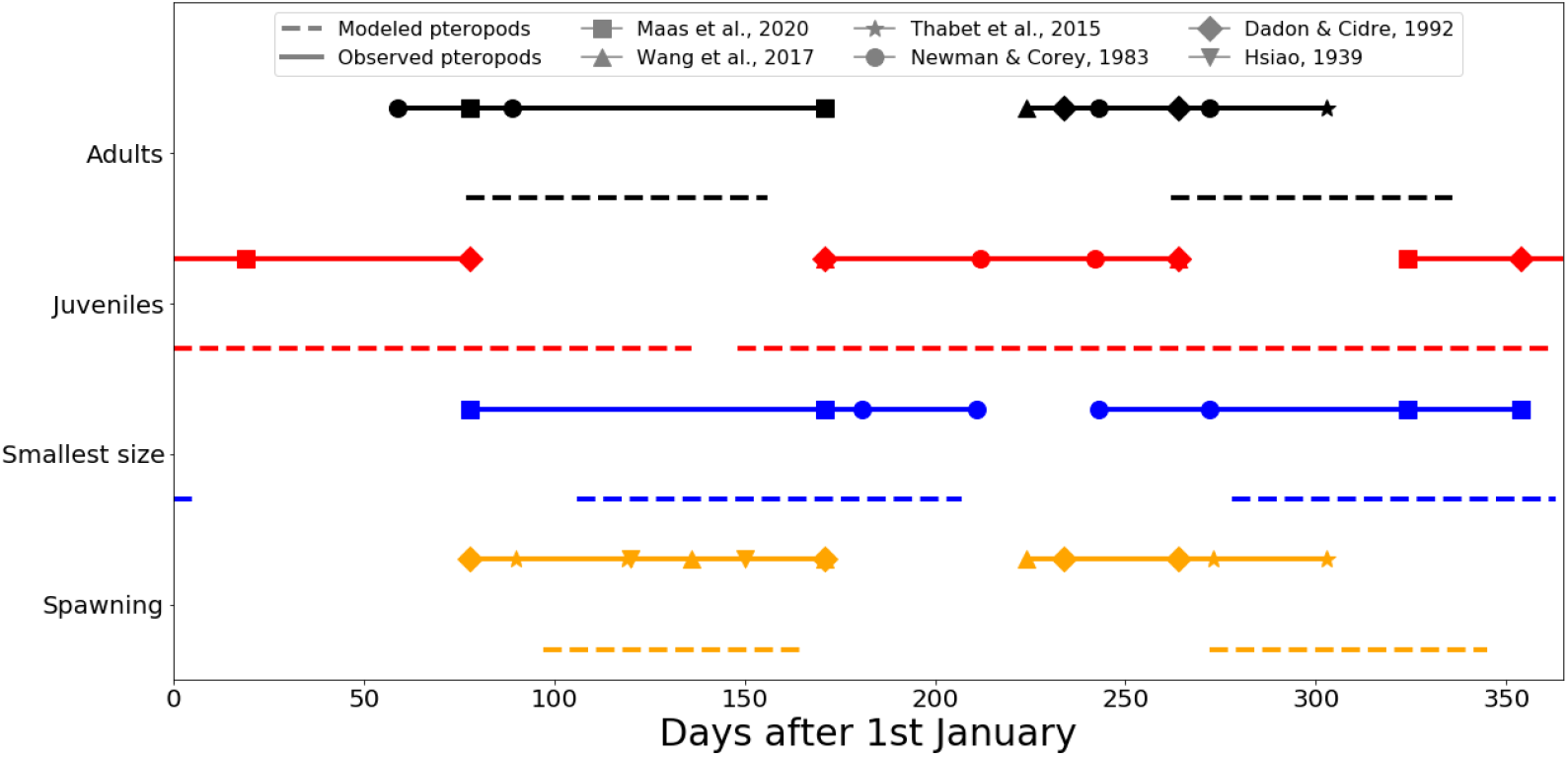
Comparison between climatological (dashed lines) and observed (solid lines) life stage occurrence throughout a year. The colors are chosen as in Figure 5, i.e. orange for eggs/spawning, blue for larvae/smallest size fraction, red for juveniles, black for adults. The symbols indicate the field observations of pteropod life stages from the following studies: upside down triangle for Hsiao (1939), diamond for Dadon and de Cidre (1992), circle for Newman and Corey (1984), star for Thabet et al. (2015), upside triangle for Wang et al. (2017), and square for Maas et al. (2020).

### 3.2. Spatio-temporal distribution of shelled pteropods

The analysis of the pteropod population dynamics neglects the spatial distribution of pteropods in the CalCS and aggregates them to a single point in space. Thus, we explore the spatio-temporal distribution of the pteropod life stages of the spring and overwintering generations throughout a year (Figure 7).

**Figure 7.**
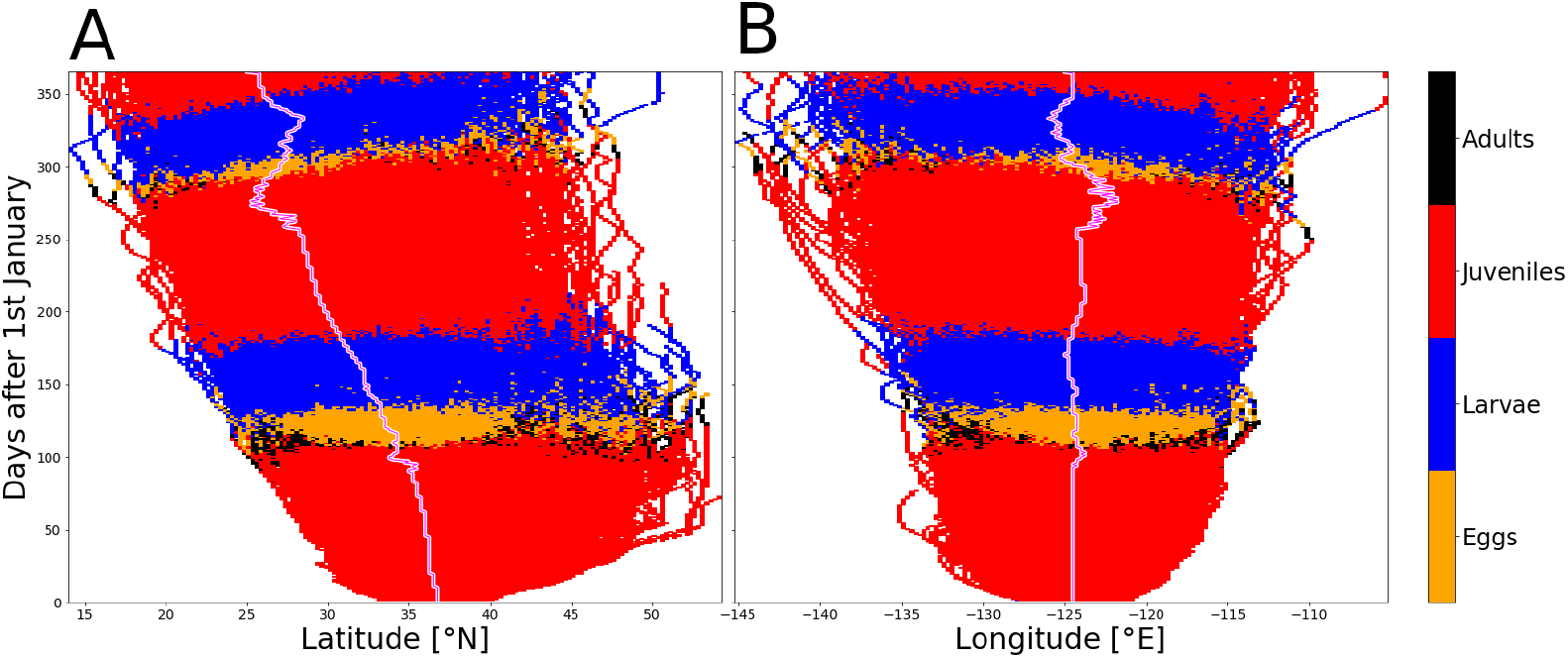
Depth integrated, dominant pteropod life stage for each simulation day and 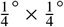 pixel in the climatology for the period between 1984 and 2019. A shows the dominant latitudinal range of the pteropod population for each simulation day. B shows the longitudinal range of the pteropod population for each simulation day. The median latitude and longitude of the population is shown in magenta with a white boarder. The colors show the four life stages eggs, larvae, juveniles, and adults in orange, blue, red, and black.

In terms of dispersal, the pteropod population is advected southwestward from its initial location (Figure 7). The initial population was located between 33°N and 40°N (median 36.75°N; Figure 7A) and between −128°E and −121°E (median −124.5°E; Figure 7B). After one year, the median latitude, and longitude of the population decreased by 11.75° (25°N; Figure 7A), and 0.5° (−125°E; Figure 7B), respectively. In addition, the range that the pteropods cover expands with time in both latitude and longitude. By the end of the simulation year, the pteropod population is located between 17°N and 44°N and between −143°E and −110°E. Overall, the juveniles are the most dominant life stage for most of the year across all latitudes (62.5%) and longitudes (63.6%; Figure 7). For roughly one quarter of the year the larval stage represents the dominant life stage across all latitudes (27.3%) and across all longitudes (27.4%). The eggs are the most dominant life stage for 7.9% (7.0%) of the year across all latitudes (longitudes). The adult life stage represent the dominant life stage only during 2.2% and 1.9% of the year across latitudes and longitudes, respectively.

In comparison to the life stage succession sequence presented above (section 3.1), we observe a similar pattern in the dominance succession across latitudes and longitudes (Figure 7). However, the adult life stage is never the dominant life stage across all latitudes or all longitudes. Starting from the 1st of January, the juvenile life stage represent the dominant life stage across all latitudes and longitudes (Figure 7). 110 (111) days after the 1st of January, eggs represents the dominant life stage until 132 (133) days after the 1st of January, when the larvae life stage becomes the most dominant life stage across all latitudes (longitudes). The larvae become the most dominant life stage between 133 (134) days after the 1st of January across all latitudes (longitudes). Between 183 and 305 (183 and 301) days after the 1st of January, the juvenile life stage becomes the most dominant life stage across all latitudes (longitudes). Afterwards, the larvae life stage dominates the simulated population until 354 (350) days after the 1st of January across all latitudes (longitudes). Finally, for the remainder of the year the juvenile life stage becomes the dominant life stage across all latitudes and longitudes.

Overall, the transitions from one life stage to the next occur roughly within the same week across latitudes during the first 180 days after the 1st of January (Figure 7A). Afterwards, the transition from one dominant life stage to next occurs later at higher latitudes than at lower ones. For instance, the transition from larvae to juveniles near the end of the year at 20°N occurs 26 days prior to the transition at 40 °N (Figure A.14). Similarly, in the first 250 days after the 1st of January, the transitions from one life stage to the next occur roughly within the same week throughout the year and across all longitudes (Figure 7B). In the second half of the year, life stages further western transition slower from one life stage to the next than in the East. Larvae located at −140 °E transition to the juvenile stage roughly 20 days later than those located at −120 °E (Figures A.15).

## 4. Discussion

Our first aim of this study was to reproduce the measured pteropod abundance signal in the temperate region (30 °N - 60 °N; section 2.10). Our shelled pteropod IBM was able to reproduce the abundance signal under time-varying environmental conditions (Figure A.22). Under these conditions, the similarity between the observed and simulated abundance is high, with a Spearman correlation coefficient of 0.96 (Figure A.22). The mismatch between simulated and observed abundance mainly occurs during early spring, where the onset of abundance increase occurs roughly 30 days later in our simulation. The delayed onset of modeled abundance increase in spring is likely linked to the narrow spawning period implemented in the IBM. However, despite the lagged onset of abundance increase in the simulation, the timing of the maximum pteropod abundance occurs within the range reported in MAREDAT, and the simulated abundance remains mostly within the observed range throughout the rest of the year (Figure A.22). Thus, on a population level, our IBM is able to accurately reproduce the pteropod abundance signal measured in the temperate region.

In addition to the population level match between simulation and observation, the second aim of this study was to provide a pteropod population life stage composition across one year that matches the life stage succession reported in the literature. In terms of longevity, our modeled pteropods have a life span of roughly seven months and eight months in the spring and overwintering generation, respectively (section 3.1). The modeled longevity is of the same order of magnitude as the longevity reported for *L. retroversa* kept under constant temperature (8° C) and food replete conditions (six months; Thabet et al., 2015), and for *L.helicina* in the temperate North Pacific (six months, and eleven months for the spring and overwintering generation, respectively; Wang et al., 2017). On the life stage level, the prolonged longevity is most pronounced in the juvenile stages. The modeled juvenile life stage has the largest difference to the reported observed duration of 60 days (section 2.7.2; Thabet et al., 2015), with a median modeled longevity of juveniles that is roughly twice as long for the spring (114 days) and overwintering (130 days; Table 3) generation. The difference between modeled and observed longevity might be explained by the difference between modeled and observed temperature and food availability that pteropods experience. Relative to the observed environmental conditions in the temperate North Pacific with temperatures ranging between 6°C and 14.5°C and chlorophyll-a concentrations between 2.5 *mg m*^−3^ and 15 *mg m*^−3^ (Wang et al., 2017), the modeled pteropods experience warmer temperatures and lower food availability in our model simulations (e.g. Figure A.16 and A.18). On one hand, the combination of warmer temperatures and food scarcity result in a slower life cycle for the spring generation, and on the other in a faster life cycle for the overwintering generation.

Detailed reports on the life stage progression or the life cycle of pteropods are scarce (Howes et al., 2014; Manno et al., 2017). However, we find that on average 86% of the modeled life stages (weighted by modeled abundance) fall within the corresponding life stage occurrence observations reported across multiple regions (Figure 6, such as in the Gulf of Maine (Hsiao, 1939; Thabet et al., 2015; Maas et al., 2020), the Bay of Fundy (Newman and Corey, 1984) the temperate North Pacific (Wang et al., 2017), or the Western South Atlantic (Dadon and de Cidre, 1992). Overall, the timing in the development of the spring generation reflects the general patterns reported in the literature (Figure 6). The main difference between modeled and observed life stages occurs for the juveniles of both generations. During winter at the beginning of the year, the simulated juveniles experience relatively cold temperatures (Figure A.16 to A.17) and likely a slower growth period (Buitenhuis et al., 2019; Thibodeau et al., 2020), which might explain the longer life stage occurrence. These lower temperatures might have extended the duration of the modeled juvenile life stage, and resulted in a progressively increasing delay in the life stage progression across the year. In relation to the life stage dominance in the model, we found that younger and smaller life stages dominate over older and larger ones in terms of abundance, which is consistent with the observed dominance of smaller life stages in the CalCS (Bednaršek and Ohman, 2015) and the temperate North Pacific (Wang et al., 2017),

The third aim of this study was to characterize the spatio-temporal distribution of pteropod life stages throughout a year. However, the comparison of model results to observations is limited by the fact that pteropods (and zooplankton in general) have a heterogeneous and patchy distribution (e.g. Wishner et al., 1988; Miller et al., 1998; Brentnall, 2003; Maas et al., 2020; Pasternak et al., 2020), where sampling pteropods across small spatial scales can yield significantly different size-specific abundances and vertical distributions in the CalCS (Bednaršek and Ohman, 2015). Nevertheless, we found key similarities between our model results and observations in terms of dispersal and life stage spatio-temporal distribution. First, we find a heterogeneous and patchy distribution in our modeled pteropod population, evidenced by the isolated paths shown in Figure 7, or as shown for seasonal maps of individual years (Figure A.20 and A.21). This aggregation into patches in our model is likely caused by eddies that are able to retain plankton within their boundaries for up to 67 days in our domain (Condie and Condie, 2016), and by our use of *super-individuals* to match the observed pteropod abundance (section 2.8). The latter results in a reduced representation of variable dispersal patterns, and further increases spatial heterogeneity in the simulation (Parry and Evans, 2008). Second, we find a predominantly southwestward trajectory of the modeled pteropod population throughout a year, which resembles the direction of the lateral export of organic matter towards the open ocean wit the California Current (Frischknecht et al., 2018). Third, our IBM simulates a dominance of the juvenile life stages in terms of presence across the year (Figure 7), and a relatively small dominance of adult life stages. This finding reflects the observation that juvenile life stages can be observed throughout the year, and the larger individuals are relatively seldom observed (Bednaršek and Ohman, 2015; Wang et al., 2017)

In summary, the abundance modeled with our shelled pteropod IBM is consistent with the measured pteropod abundance signal in the temperate region of the Northern Hemisphere, and its application to the CalCS results in a realistic perspective of the life stage progression for the simulated years. This is evidenced by the match between the simulated life stage progression and the reported occurrence of life stages across several regions (Figure 6), such as the major spawning events in spring and fall (Hsiao, 1939; Dadon and de Cidre, 1992; Thabet et al., 2015), the subsequent die-off events (Dadon and de Cidre, 1992; Maas et al., 2020), the overall dominance of earlier life stages in terms of abundance and dominant presence of juvenile life stages throughout a year (Newman and Corey, 1984; Dadon and de Cidre, 1992; Wang et al., 2017).

### 4.1. Caveats and limitations

Despite the large amount of information on the biology of shelled pteropods, information on some key processes remains scarce, such as life stage specific mortality (Lischka et al., 2011; Bednaršek et al., 2012b, 2016). As shown by our sensitivity analysis, changes in the mortality rates mainly affect the modeled abundances but not the seasonality (Table A.6). In terms of abundance changes linked to mortality rates uncertainty, the shelled pteropod IBM uses *super-individuals* to match the observed pteropod abundance. Thus, we do not expect the life stage succession patterns to be affected by this uncertainty, and the decrease/increase in pteropod abundance due to the uncertainties in mortality rates could be lessened by adjusting the number of individuals within each *super-individual*.

In terms of the spatio-temporal distribution of pteropods, a potential source of uncertainty is our choice in restarting each simulation year with the same initial population (section 2.10), and the use of *super-individuals* itself. The former led to a simulation with reduced variability in the life stage composition of the pteropod population at the beginning of the simulation and a higher life stage variability near the end of the simulation. However, this approach emphasizes the role of environmental conditions, such as temperature (Figure A.16) or food availability (Figure A.19) on the population life stage progression (see also Dorman et al., 2011, 2015a,b). With respect to the use of *super-individuals*, this approach was necessary to improve the model speed and memory use (Scheffer et al., 1995), at the cost of reduced variable dispersal and enhanced patchiness (Parry and Evans, 2008). However, unlike in our pteropod IBM, the reduced dispersal is especially important if density-dependent functions and interactive individuals are used (Parry and Evans, 2008). In addition, the number of individuals per *super-individual* in our IBM is roughly within the recommended order of magnitude (ten; Parry and Evans, 2008).

Other potential caveats relate to the responses of pteropod to environmental conditions. Our model does not incorporate a direct pteropod response to some key factors such as oxygen (Gilmer, 1974; McLaren, 1963; Hauss et al., 2016) or predation (e.g. by fish, or shell-less pteropods; Ohman, 1990; Peijnenburg et al., 2019). However, additional experimental and field work is needed prior to the incorporation of these direct responses, due to the relatively high uncertainty in the influence of environmental stressors on pteropods and missing parameterisations (Lischka et al., 2011; Lischka and Riebesell, 2016; Pasternak et al., 2020).

## 5. Conclusion

To the best of our knowledge, this is the first IBM to simulate the life cycle of shelled pteropods with a spring and overwintering generation and four life stages. In the model we use available information on, and parameterisations of the biology of *L. helicina* and *L. retroversa*, and link the characteristics of the individuals (e.g. growth, the onset of vertical migration, mortality, development, response to environmental conditions) to their size and the environmental conditions that they experience throughout their life. Our model successfully captures the measured pteropod abundance signal of the regions between 30°N and 60°N, the life stage duration, and succession sequence reported in the literature. As shown in this study, our model can be used to characterize the spatio-temporal distribution of pteropod life stages in a region (e.g. CalCS).

Overall, our IBM is particularly relevant for quantifying ongoing and future effects of climate change, which may manifest as changes in pteropod abundance, size, or growth (e.g. ocean acidification; Bednaršek et al., 2017b). Our implementation of size as a key determinant in the behaviour and development of pteropods (see Litchman et al., 2013), can be used to illustrate possible changes in the phenology and life stage composition of a pteropod population throughout a year (Figure 5), and keep track of the location and environmental conditions that pteropods experience (e.g. food availability; Figure A.18). In addition, the inclusion of mechanistic responses to environmental conditions may allow our IBM to capture the effects of environmental stressors on populations as a whole, or on individuals (Marshall et al., 2017), as well as the effects of multiple stressors on pteropods (Doo et al., 2020) in the context of climate change. The latter may provide a more detailed picture of the effects of climate change on marine ecosystems, where pteropod shell dissolution already today is used to illustrate a concrete and tangible marine ecosystem response to climate change (Manno et al., 2017; Bednaršek et al., 2019).

## Acknowledgments

We thank N. Gruber and the Environmental Physics group for analytical and visualization advice.

## Author contributions

M.V. and U.H.E. conceptualized the project. U.H.E. and M.V. developed the model. U.H.E. prepared the data, performed the analyses and produced the visualizations. U.H.E. wrote the first draft with substantial inputs by M.V. Both authors interpreted the results. All authors reviewed the manuscript. M.V. supervised the project.

## Data and materials availability

The code for the project is freely available upon request.

## Appendix A. Appendix

### Appendix A.1. Shell growth

**Figure A.8.**
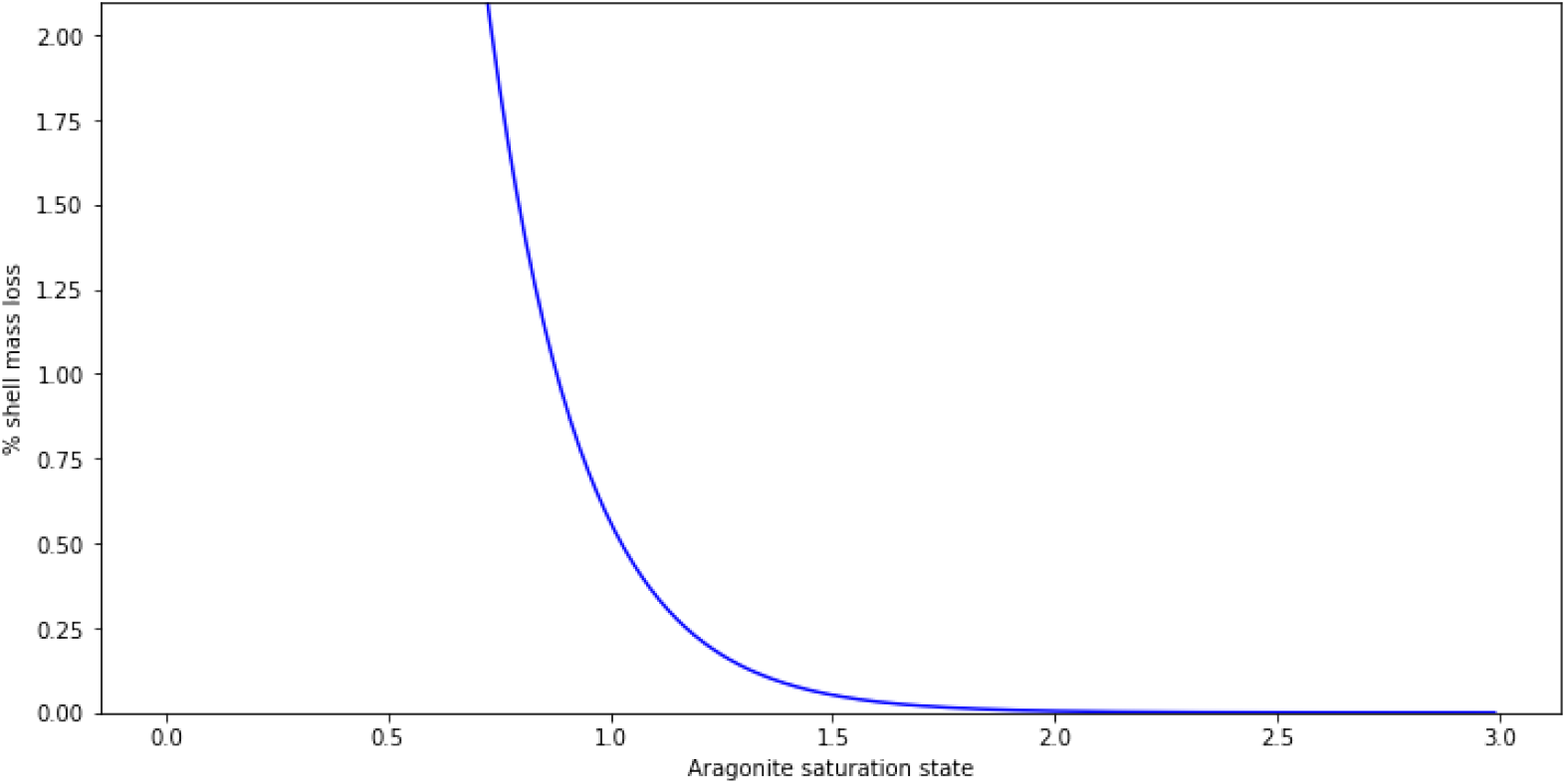
Percentage of shell mass loss as a function of aragonite saturation state. The equation was taken from Bednaršek et al. (2014).

### Appendix A.2. Mortality increase due to exposure to corrosive waters

We calculate the linear increase in mortality coefficient (M) of modeled pteropods as a function of aragonite saturation state (Ω_*arag*_; Figure A.9). To do this, we use the experimental observations from Lischka et al. (2011), for the mortality of *L. helicina* under different combinations of Ω_*arag*_ and temperatures that were reported to be within the natural range of the pteropod species. This resulted in the relationship *M* = −0.06 · Ω_*arag*_ + 0.50.

**Figure A.9.**
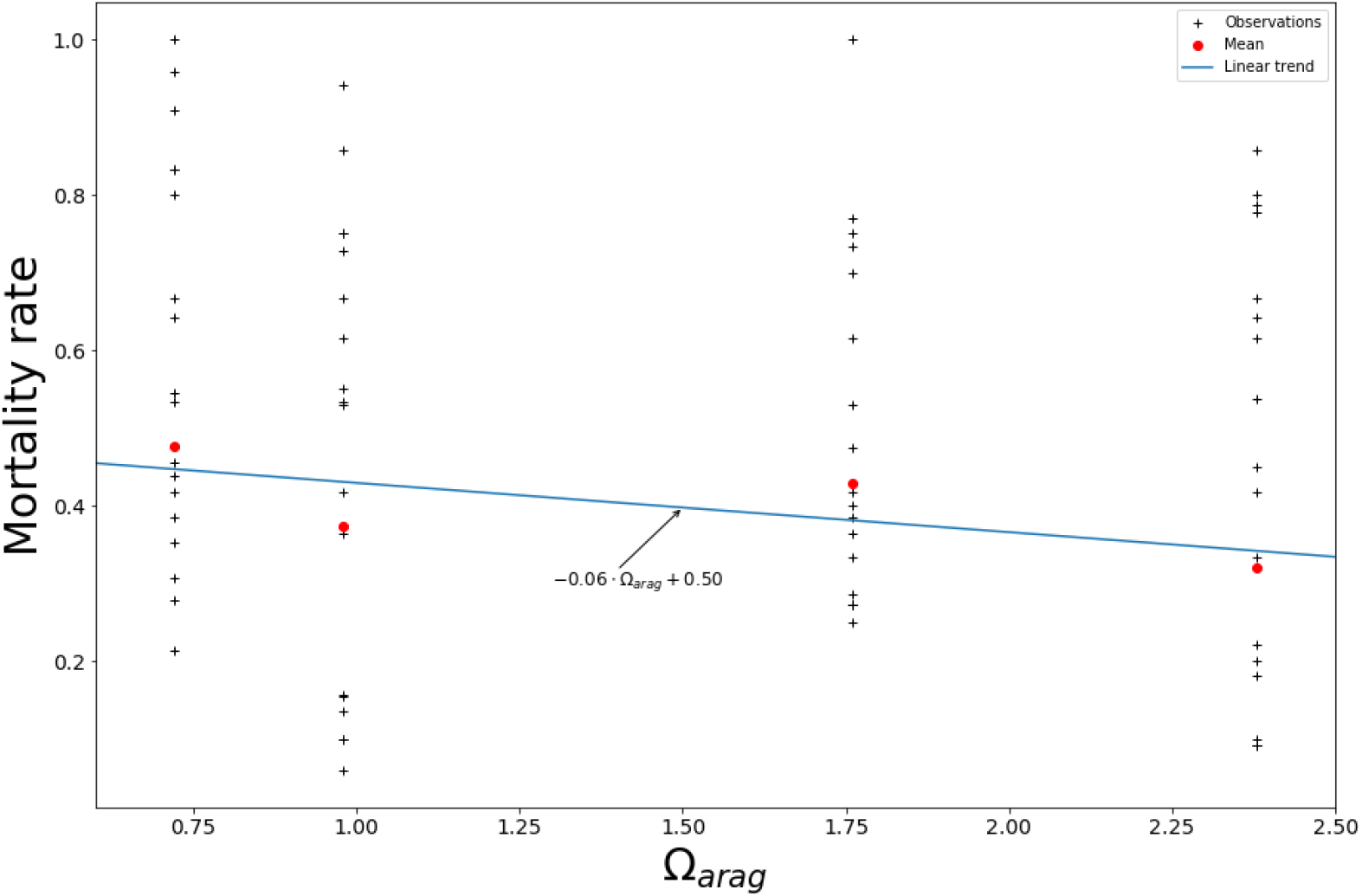
Mortality rate as a function of aragonite saturation state (Ω_*arag*_) based on the observations presented in Lischka et al. (2011). The linear trend was calculated based on all observations where the experimental temperatures were reported to be within the natural range of *L. helicina*, i.e. observations done at 3°C and 5.5°C. Black crosses denote the individual observations, the red dots the mean, and the blue line the linear trend.

### Appendix A.3. Model functions and parameters

**Table A.4.**
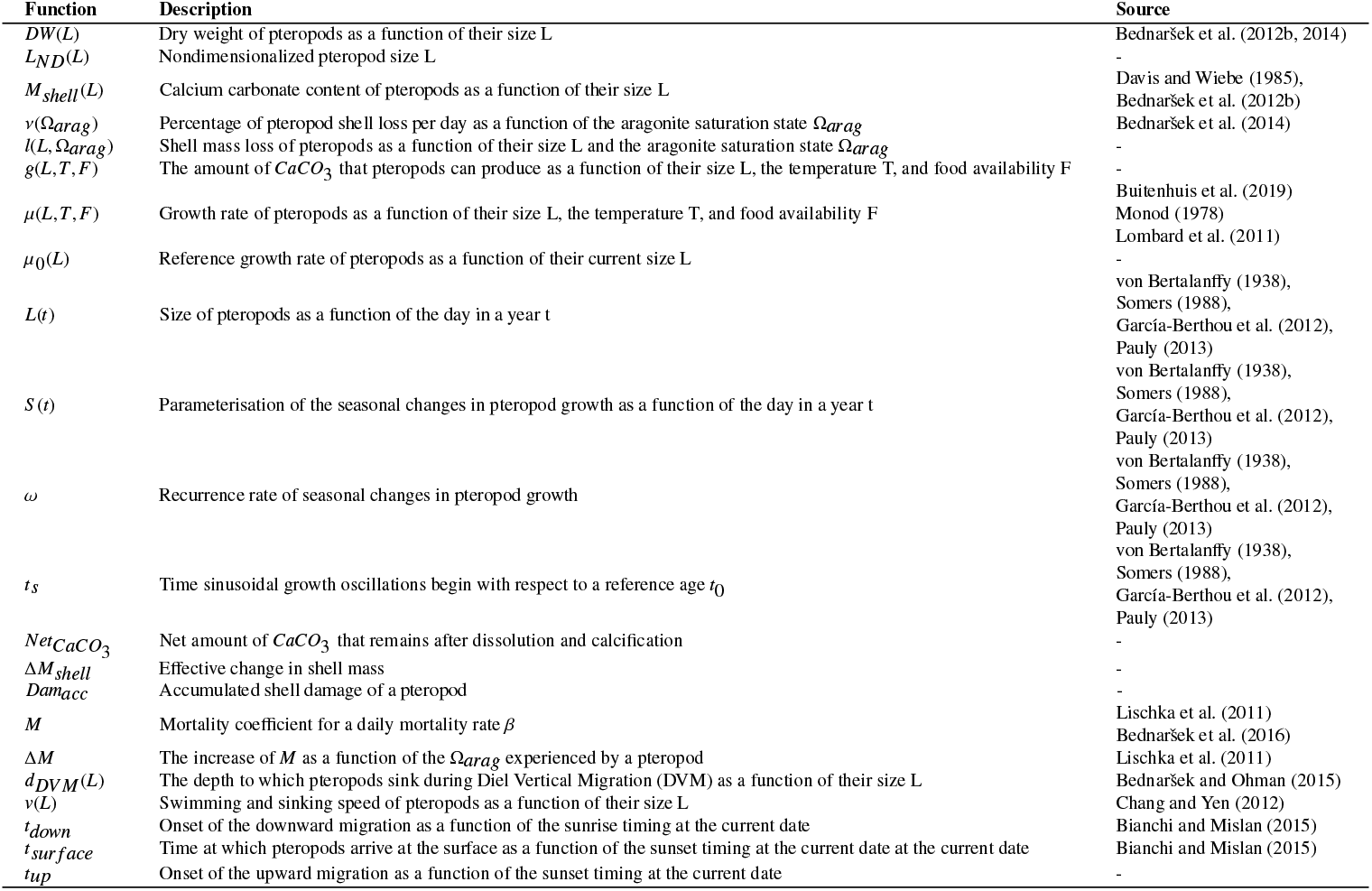
Functions used in our shelled pteropod Individual-based model.

**Table A.5.**
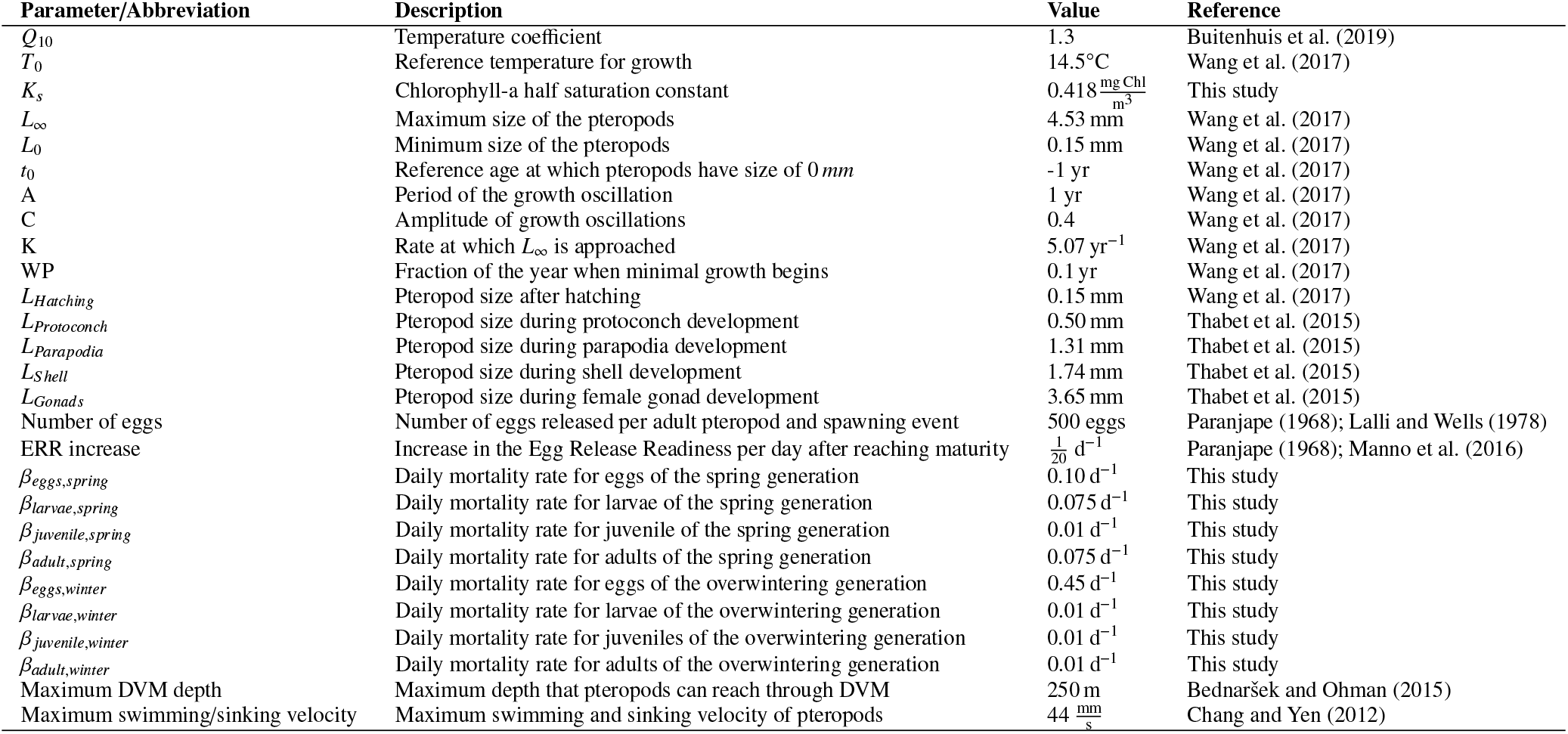
Shelled pteropod Individual-based model parameters chosen for this study.

### Appendix A.4. Model tuning

**Figure A.10.**
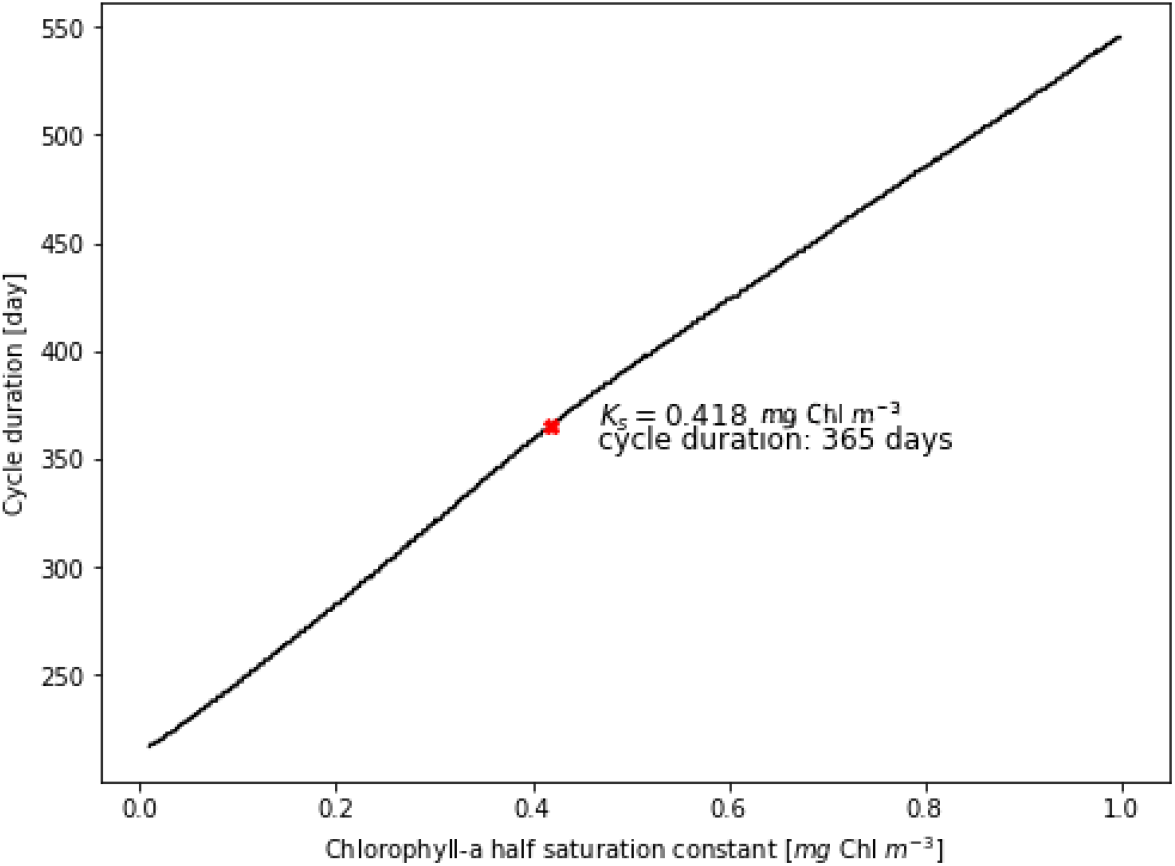
Time span between a pteropod that hatches in spring (30th April) and the day that the next generation releases eggs as a function of the chlorophyll-a half saturation constant for food limited growth. The red dot shows the half saturation constant where the IBM simulates the reported two generation per year cycle for temperate shelled pteropods (Lalli and Gilmer, 1989; Dadon and de Cidre, 1992; Wang et al., 2017; Maas et al., 2020).

**Figure A.11.**
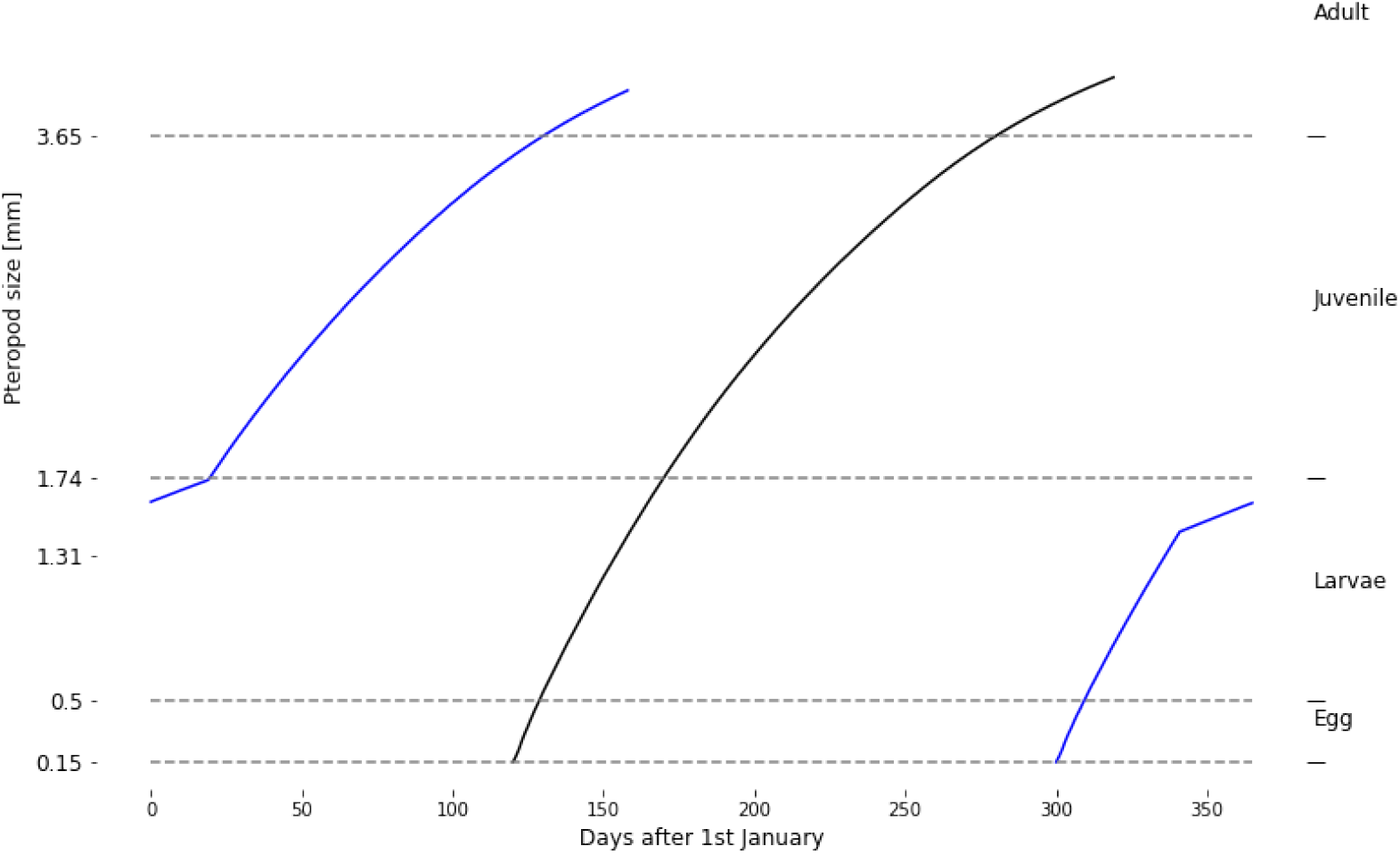
Size of pteropods parameterised as a function of temperature and chlorophyll-a concentration under idealized temperature and food scarcity during winter. In black is the size of pteropods of the spring generation. In blue is the size of pteropods of the overwintering generation.

### Appendix A.5. Sensitivity analysis

**Table A.6.** Outcome of the sensitivity analysis for the life stage and generation specific mortality rates, the number of eggs released per adult pteropod and spawning event, and the daily increase in the Egg Release Readiness (ERR) index after reaching maturity. The sensitivity was quantified using the Manhattan distance (*L*1) and the Pearson correlation (*r*) between the modeled pteropod abundance and the pteropod abundance reported in MAREDAT (Bednaršek et al., 2012a) between 30°N and 60°N. The sensitivity experiments were run under idealized conditions (section 2.5) with chosen parameters values increased/decreased by 10% individually. Under idealized conditions our IBM has *L*1 = 15.5 and *r* = 0.99.

| Parameter | Value | L1 | r |
| --- | --- | --- | --- |
| $\beta_{egg, spring}$ | 0.11 d<sup>-1</sup> | 20.0 | 0.94 |
|  | 0.09 d<sup>-1</sup> | 19.4 | 0.98 |
| $\beta_{larvae, spring}$ | 0.0825 d<sup>-1</sup> | 24.4 | 0.97 |
|  | 0.0675 d<sup>-1</sup> | 15.5 | 0.98 |
| $\beta_{juvenile, spring}$ | 0.011 d<sup>-1</sup> | 22.3 | 0.95 |
|  | 0.009 d<sup>-1</sup> | 16.6 | 0.98 |
| $\beta_{adult, spring}$ | 0.0825 d<sup>-1</sup> | 21.7 | 0.95 |
|  | 0.0675 d<sup>-1</sup> | 21.2 | 0.97 |
| $\beta_{egg, winter}$ | 0.495 d<sup>-1</sup> | 25.3 | 0.95 |
|  | 0.405 d<sup>-1</sup> | 16.2 | 0.98 |
| $\beta_{larvae, winter}$ | 0.011 d<sup>-1</sup> | 21.4 | 0.97 |
|  | 0.009 d<sup>-1</sup> | 20.4 | 0.97 |
| $\beta_{juvenile, winter}$ | 0.011 d<sup>-1</sup> | 17.8 | 0.98 |
|  | 0.009 d<sup>-1</sup> | 19.2 | 0.97 |
| $\beta_{adult, winter}$ | 0.011 d<sup>-1</sup> | 18.3 | 0.98 |
|  | 0.009 d<sup>-1</sup> | 19.4 | 0.98 |
| Number of eggs | 550 eggs | 15.8 | 0.99 |
|  | 450 eggs | 27.1 | 0.87 |
| Increase in ERR | 0.04 | 20.7 | 0.99 |
|  | 0.07 | 22.6 | 0.98 |

**Figure A.12.**
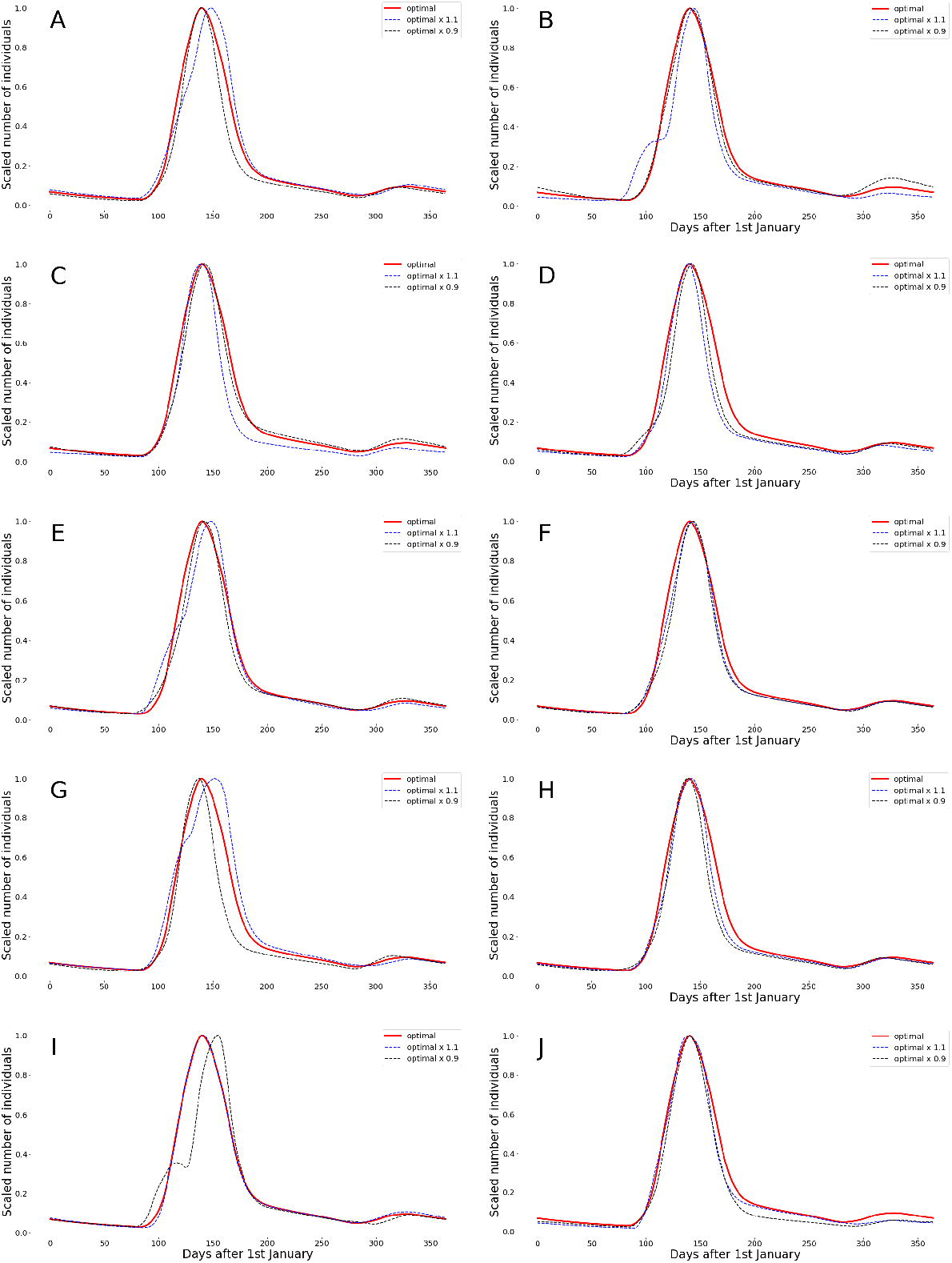
Comparison of modeled pteropod abundance after increasing/decreasing the life stage and generation specific mortality rates for eggs of the (A) spring and (B) overwintering generation, for larvae of the (C) spring and (D) overwintering generation, for juveniles of the (E) spring and (F) overwintering generation, for adults of the (G) spring and (H) overwintering generation, the (I) number of eggs released per adult pteropod and spawning event, and the (J) daily increase in the Egg Release Readiness (ERR) index after reaching maturity. The red curve shows the modeled pteropod abundance without changing the aforementioned parameters, the blue/black curve shows the abundance after increasing/decreasing the parameter by 10%. Each modeled abundance was scaled to the interval between zero and one for comparison.

### Appendix A.6. Seeding region

The pteropods were seeded randomly with repetition within the region shown in Figure A.13 during the initialization. This region is between 33°N and 48°N, and between 128°W and 121°W. This seeding approach allowed us to maintain the pteropod population in the region of interest, and characterize their horizontal dispersion throughout a year.

**Figure A.13.**
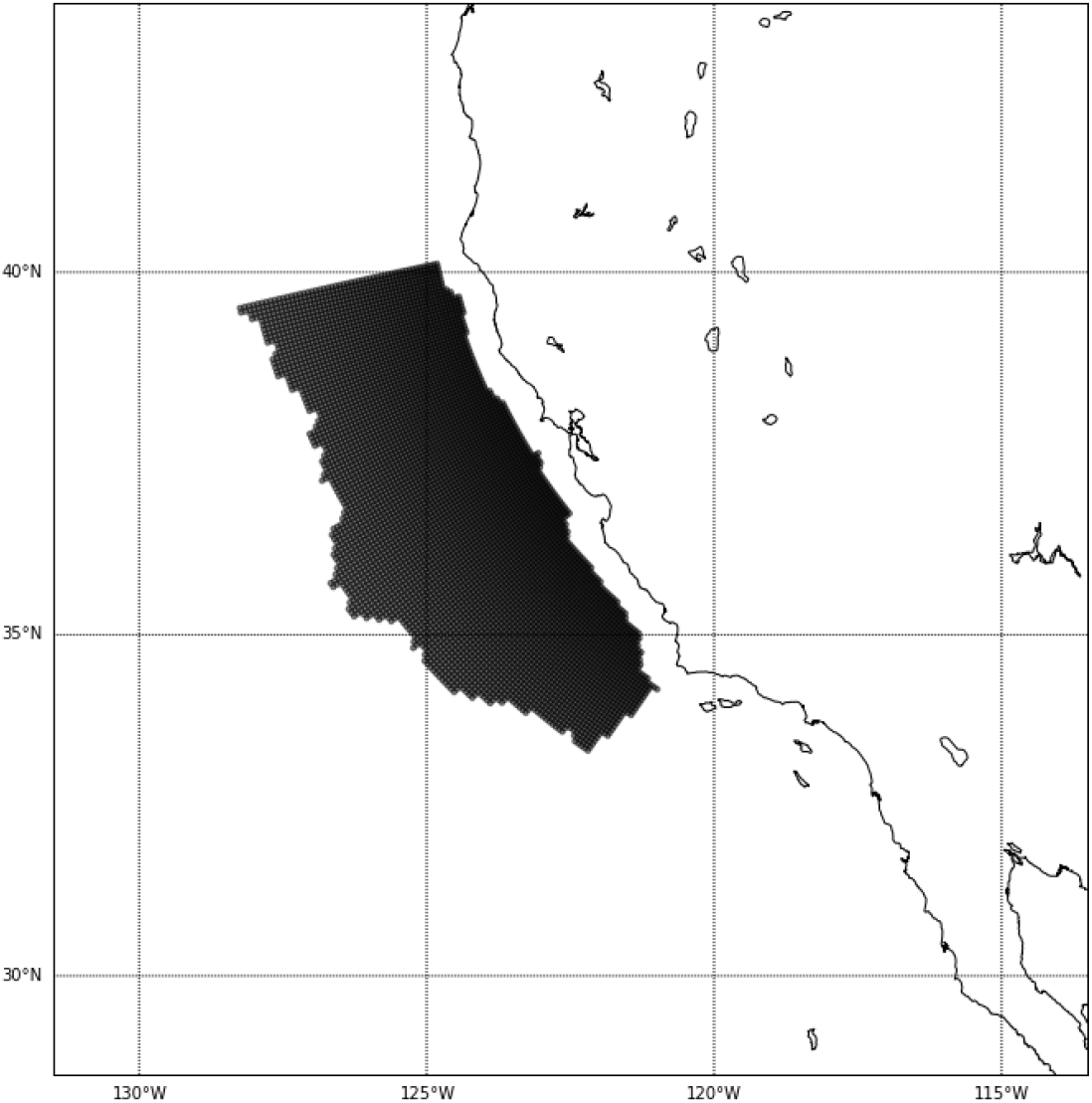
Area in the California Current System where the pteropods were seeded.

### Appendix A.7. Spatio-temporal life stage succession patterns

**Figure A.14.**
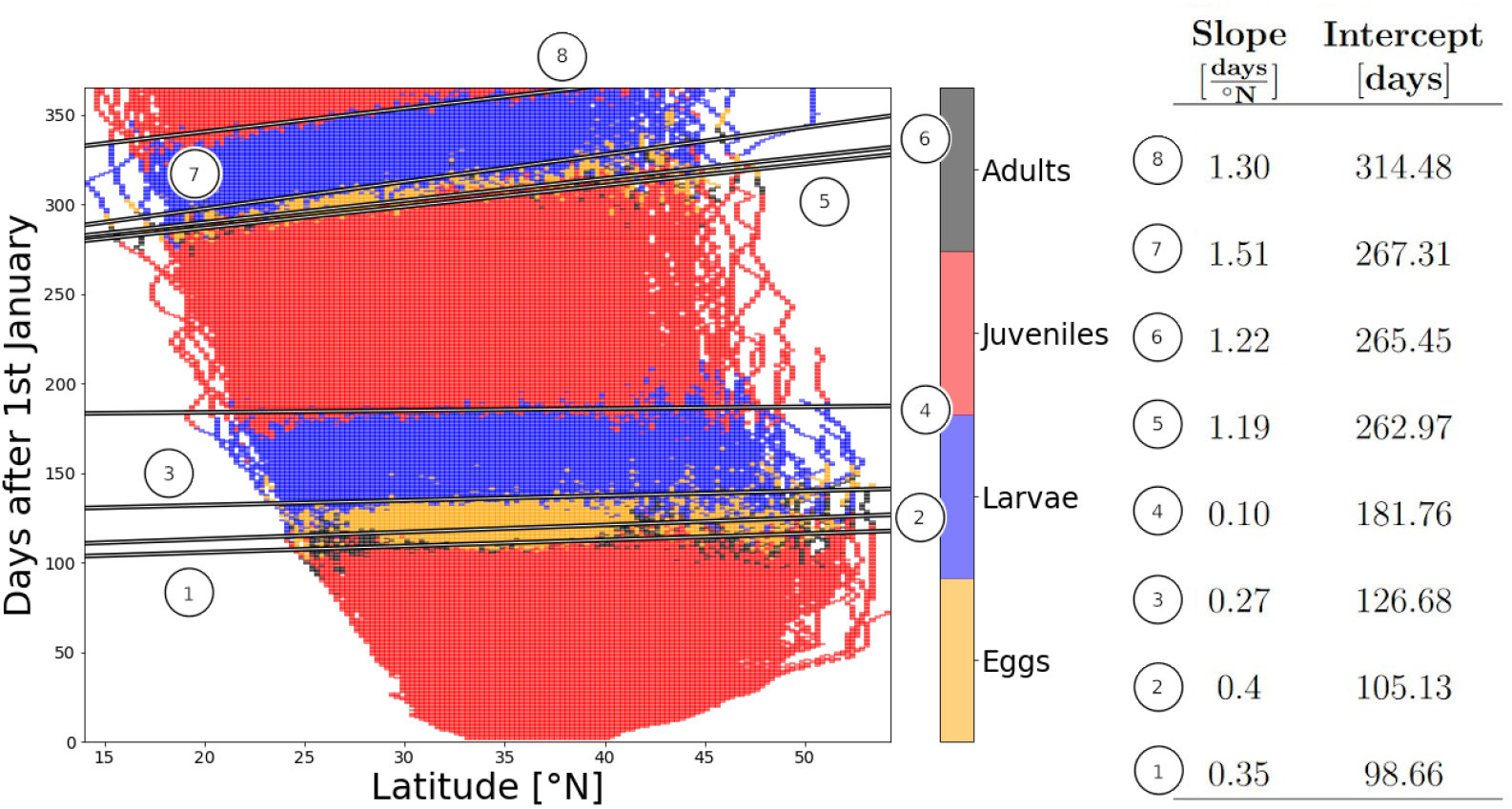
Life stage succession timing as a function of latitude. The linear trend was calculated for each succession from one life stage to the next life stage shown by the depth integrated, dominant pteropod life stage for each simulation day and 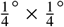 pixel in the climatology simulation. The colors show the four life stages eggs, larvae, juveniles, and adults in orange, blue, red, and black, respectively.

**Figure A.15.**
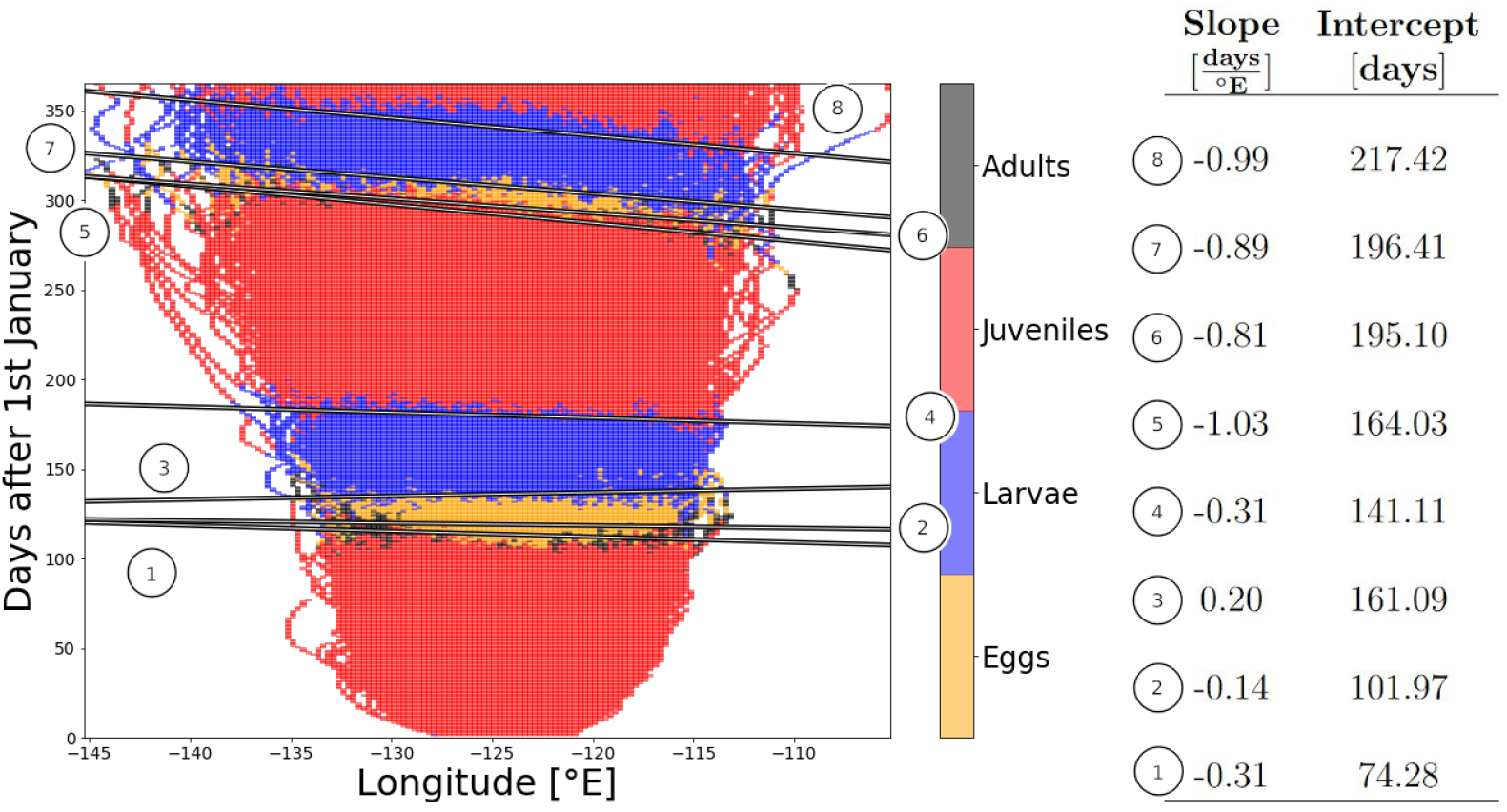
Life stage succession timing as a function of longitude. The linear trend was calculated for each succession from one life stage to the next life stage shown by the depth integrated, dominant pteropod life stage for each simulation day and 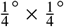 pixel in the climatology simulation. The colors show the four life stages eggs, larvae, juveniles, and adults in orange, blue, red, and black, respectively.

**Figure A.16.**
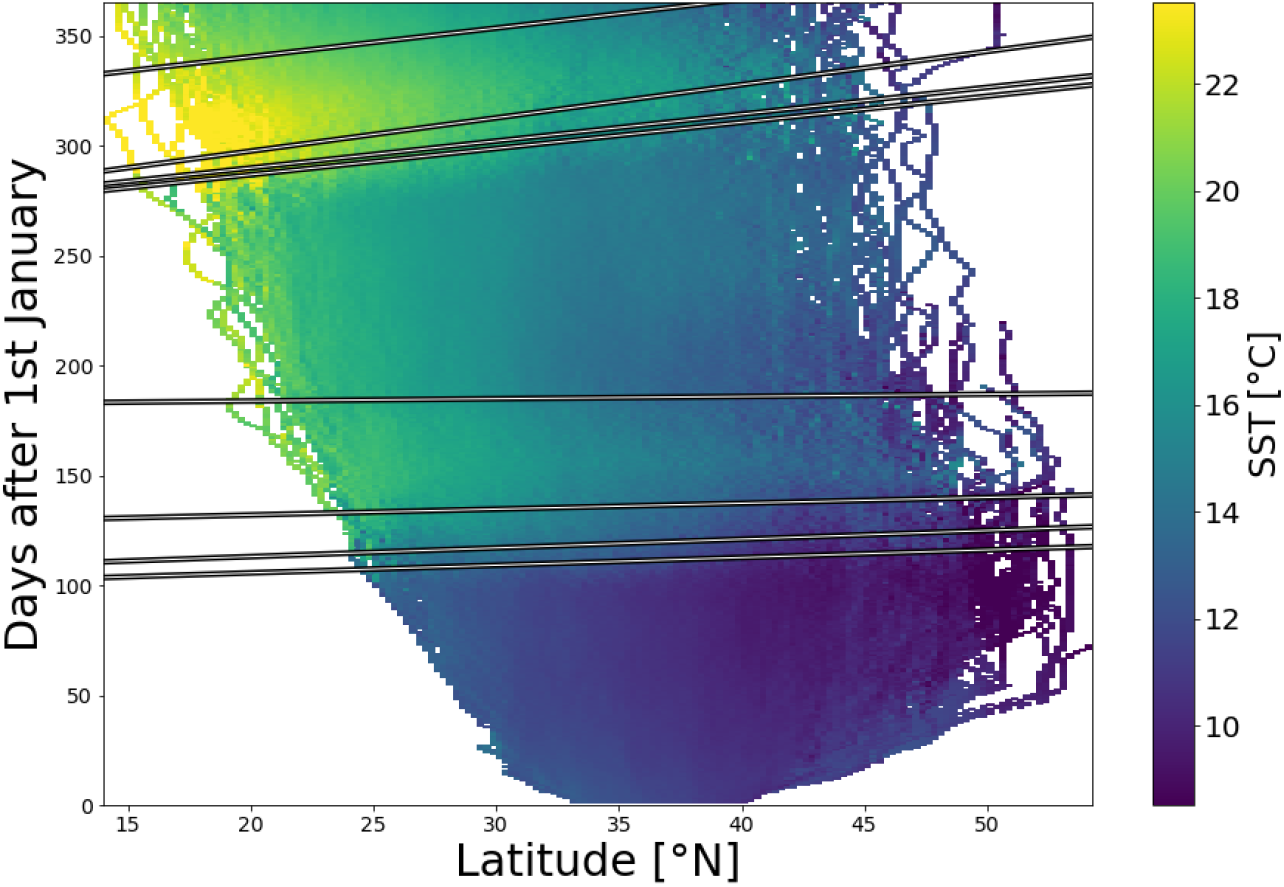
Mean temperature experienced by pteropods as a function of latitude and time. The linear trend was calculated for each succession from one life stage to the next life stage shown by the depth integrated, dominant pteropod life stage for each simulation day and 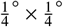 pixel in the climatology simulation.

**Figure A.17.**
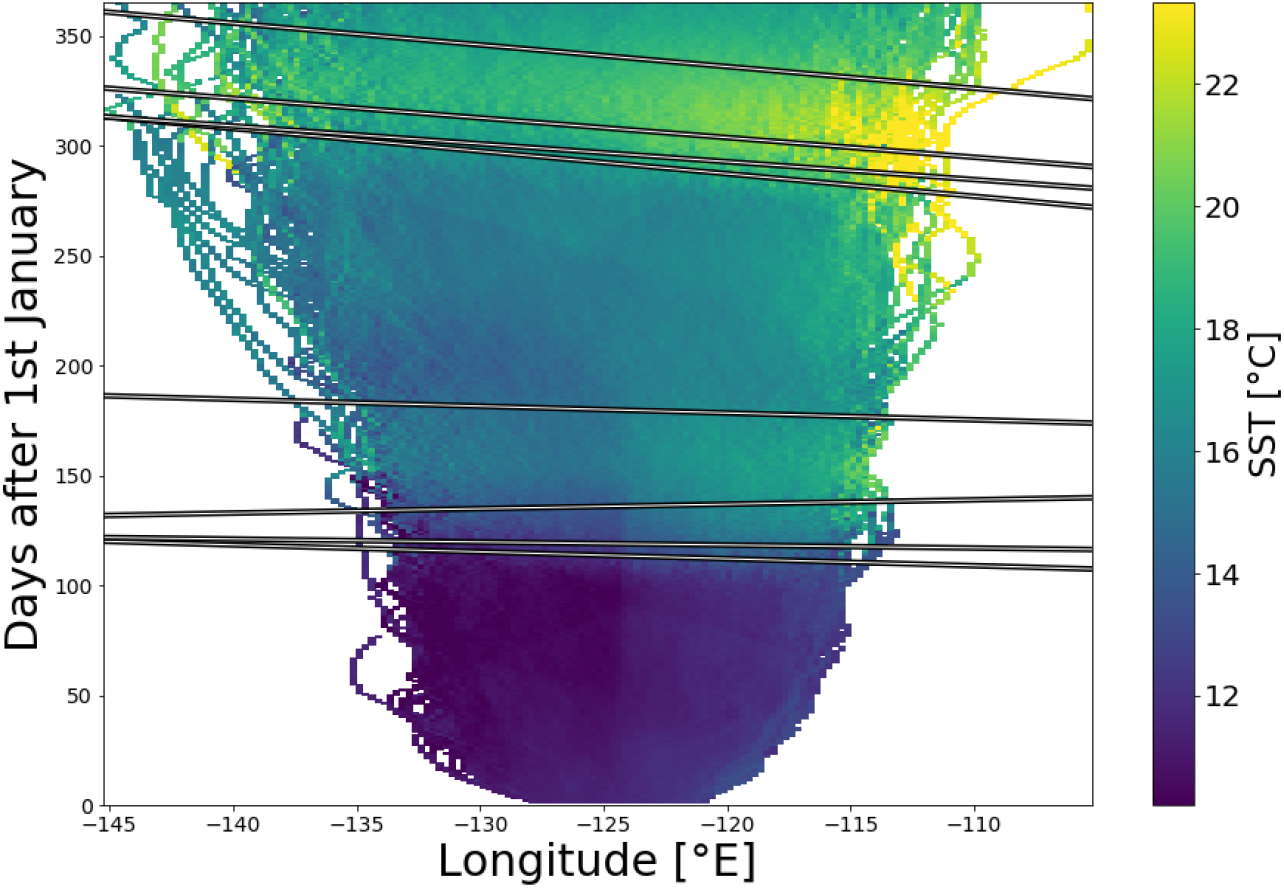
Mean temperature experienced by pteropods as a function of longitude and time. The linear trend was calculated for each succession from one life stage to the next life stage shown by the depth integrated, dominant pteropod life stage for each simulation day and 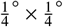 pixel in the climatology simulation.

**Figure A.18.**
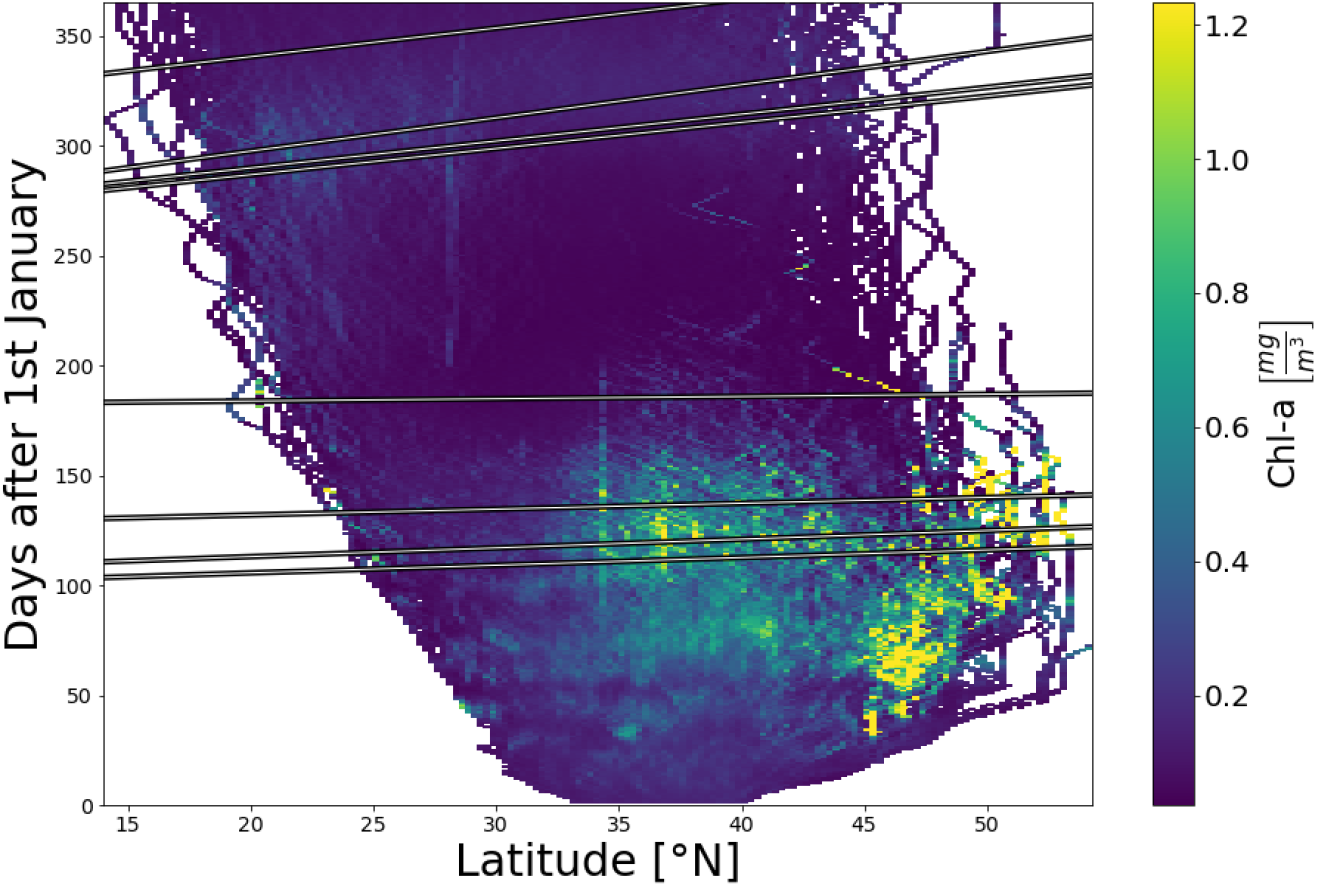
Mean cumulative chlorophyll-a concentration experienced by pteropods as a function of latitude and time. The linear trends were calculated for each succession from one life stage to the next life stage shown by the depth integrated, dominant pteropod life stage for each simulation day and 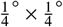 pixel in the climatology simulation.

**Figure A.19.**
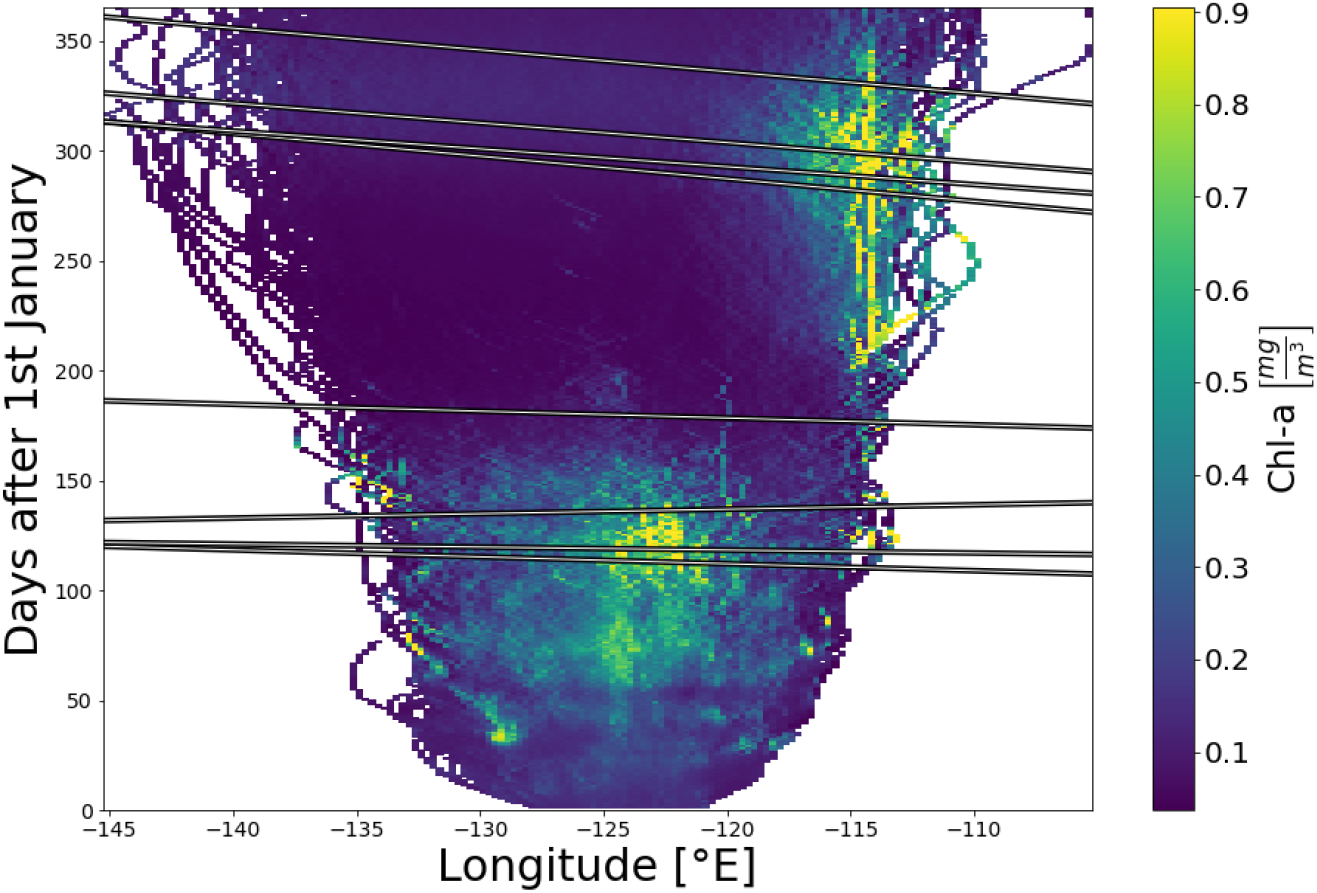
Mean cumulative chlorophyll-a concentration experienced by pteropods as a function of longitude and time. The linear trends were calculated for each succession from one life stage to the next life stage shown by the depth integrated, dominant pteropod life stage for each simulation day and 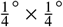 pixel in the climatology simulation.

### Appendix A.8. Seasonal distribution of pteropod life stages

**Figure A.20.**
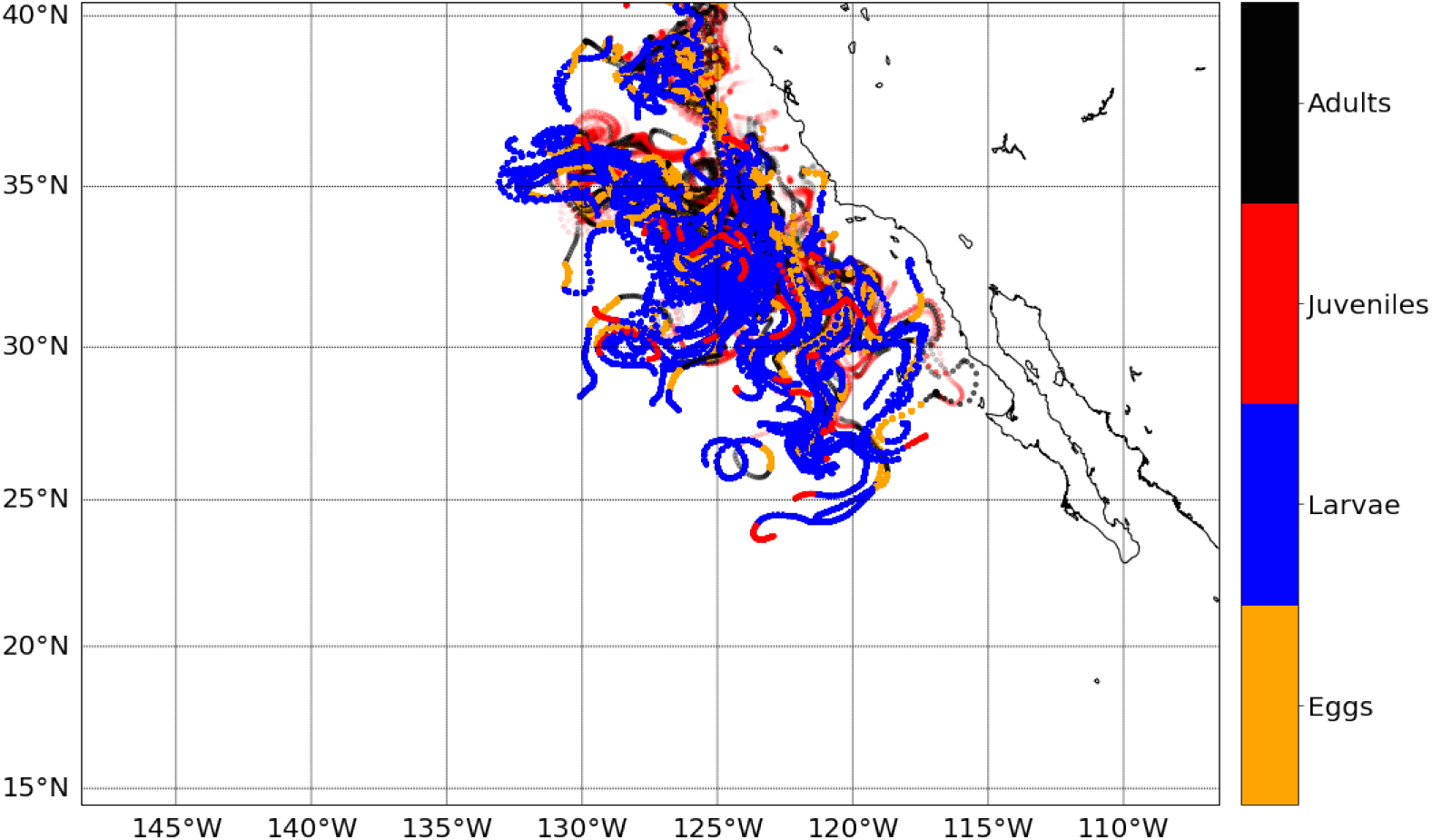
Simulated distribution and trajectories of pteropod life stages during spring of the year 2000. The trajectories are shown by the transparency. The transparency decreases with each day after the beginning of the season and turns opaque at the end of the season. The colors show the four life stages eggs, larvae, juveniles, and adults in orange, blue, red, and black, respectively.

**Figure A.21.**
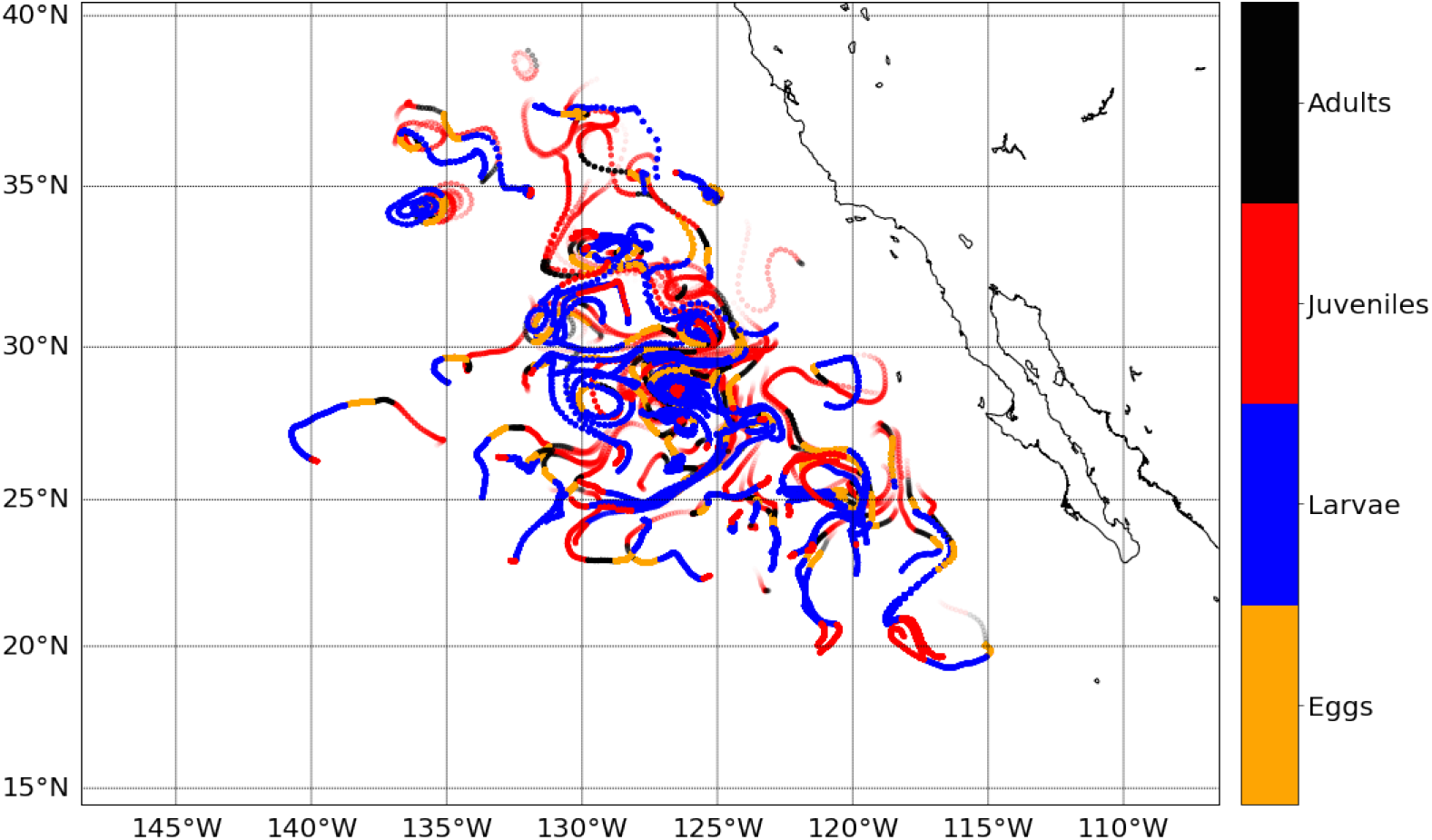
Simulated distribution and trajectories of pteropod life stages during fall of the year 2000. The trajectories are shown by the transparency. The transparency decreases with each day after the beginning of the season and turns opaque at the end of the season. The colors show the four life stages eggs, larvae, juveniles, and adults in orange, blue, red, and black, respectively.

### Appendix A.9. Comparison of modeled abundance and MAREDAT observations under non-idealized conditions

**Figure A.22.**
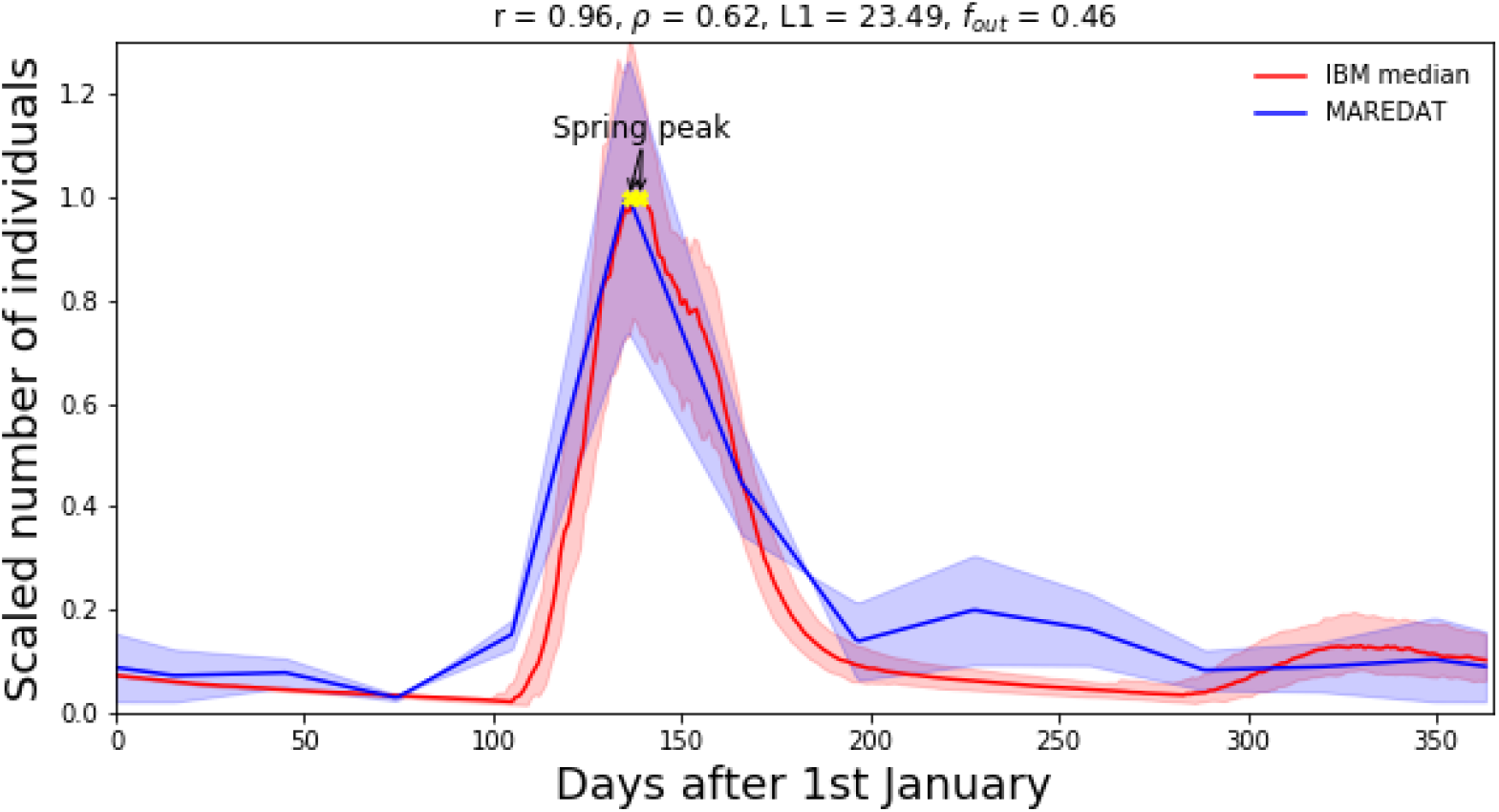
Abundance climatology (median) and 90th percentile confidence interval of simulated pteropods (red), and the daily abundance data and standard deviation of pteropods reported in the MAREDAT initiative (blue; Bednaršek et al., 2012a). The MAREDAT daily data was interpolated from monthly averaged and seasonally corrected abundance data found between 30°and 60°latitude in the Northern Hemisphere. The abundance data was scaled to the interval between zero and one. The spring pteropod abundance peak is shown in yellow. The Pearson correlation coefficient (*r*), Spearman correlation coefficient (*ρ*), the Manhattan distance (*L*1), and the proportion of modeled abundance outside of the measured abundance range (*f_out_*) between the MAREDAT data and the climatology are given at the top of the figure.

